# Small molecules inhibit SARS-COV-2 induced aberrant inflammation and viral replication in mice by targeting S100A8/A9-TLR4 axis

**DOI:** 10.1101/2020.09.09.288704

**Authors:** Qirui Guo, Yingchi Zhao, Junhong Li, Jiangning Liu, Xuefei Guo, Zeming Zhang, Lili Cao, Yujie Luo, Linlin Bao, Xiao Wang, Xuemei Wei, Wei Deng, Luoying Chen, Hua Zhu, Ran Gao, Chuan Qin, Xiangxi Wang, Fuping You

## Abstract

The SARS-CoV-2 pandemic poses an unprecedented public health crisis. Accumulating evidences suggest that SARS-CoV-2 infection causes dysregulation of immune system. However, the unique signature of early immune responses remains elusive. We characterized the transcriptome of rhesus macaques and mice infected with SARS-CoV-2. Alarmin S100A8 was robustly induced by SARS-CoV-2 in animal models as well as in COVID-19 patients. Paquinimod, a specific inhibitor of S100A8/A9, could reduce inflammatory response and rescue the pneumonia with substantial reduction of viral titers in SASR-CoV-2 infected animals. Remarkably, Paquinimod treatment resulted in 100% survival of mice in a lethal model of mouse coronavirus (MHV) infection. A novel group of neutrophils that contributed to the uncontrolled inflammation and onset of COVID-19 were dramatically induced by coronavirus infections. Paquinimod treatment could reduce these neutrophils and regain antiviral responses, unveiling key roles of S100A8/A9 and noncanonical neutrophils in the pathogenesis of COVID-19, highlighting new opportunities for therapeutic intervention.

## Introduction

The ongoing Corona Virus Disease 2019 (COVID-19) caused by severe acute respiratory syndrome coronavirus-2 (SARS-CoV-2) has resulted in unprecedented public health crises, requiring a deep understanding of the pathogenesis and developments of effective COVID-19 therapeutics (Wu et al., 2020b; Zhu et al., 2020). Innate immunity is an important arm of the mammalian immune system, which serves as the first line of host defense against pathogens. Most of the cells of the body harbor the protective machinery of the innate immunity and can recognize foreign invading viruses (Akira et al., 2006). The innate immune system recognizes microorganisms via the pattern-recognition receptors (PRRs) and upon detection of invasion by pathogens, and PRRs activate downstream signaling pathways leading to the expression of various cytokines and immune-related genes for clearing the pathogens including bacteria, viruses and others (Akira et al., 2006). With regards to SARS-CoV-2 infection, an overaggressive immune response has been noted which causes immunopathology (Huang et al., 2020; Zhang et al., 2020). In addition, T cell exhaustion or dysfunction has also been observed (Diao et al., 2020; Zheng et al., 2020a; Zheng et al., 2020b). Besides, some studies suggest that there may be a unique immune response evoked by coronaviruses (Blanco-Melo et al., 2020). However, the nature of these responses elicited by the virus remains poorly understood.

Accumulating evidences suggest that the neutrophil count is significantly increased in COVID-19 patients with severe symptoms (Kuri-Cervantes et al., 2020; Liao et al., 2020; Tan et al., 2020; Wu et al., 2020a). It is believed that neutrophils migrate from the circulating blood to infected tissues in response to inflammatory stimuli, where they protect the host by phagocytosing, killing and digesting bacterial and fungal pathogens (Nauseef and Borregaard, 2014; Nicolas-Avila et al., 2017). The role of such a response in host defense against viral infection has not been clearly characterized. A recent study observed a new subpopulation of neutrophils in COVID-19 patients, which have been named developing neutrophils because they lack canonical neutrophil markers like CXCR2 and FCGR3B (Wilk et al., 2020). However, it is still not clear how this type of neutrophil is activated. Moreover, the precise function of these cells is also unknown.

Alarmins are endogenous, chemotactic and immune activating proteins/peptides which are released as a result of cell injury or death, degranulation, or in response to infection. They mediate the relay of intercellular defense signals by interacting with chemotactic and pattern-recognition receptors (PRRs) to activate immune cells in host defense (Oppenheim and Yang, 2005; Yang et al., 2017). Currently, the major categories of alarmins include defensins, high-mobility group (HMG) proteins, interleukins (ILs), heat shock proteins (HSPs), S100 proteins, uric acid, hepatoma derived growth factor (HDGF), eosinophil-derived neurotoxin (EDN), and cathelin-related antimicrobial peptide (CRAMP) (Giri et al., 2016; Yang et al., 2017). In response to microbial infection, alarmins are released to initiate and amplify innate/inflammatory immune responses, which involve the activation of resident leukocytes (e.g. macrophages, dendritic cells, mast cells, etc.), production of inflammatory mediators (cytokines, chemokines, and lipid metabolites), recruitment of neutrophils and monocytes/macrophages for the purpose of eliminating invading microorganisms and clearing injured tissues (Bianchi, 2007; Chen and Nunez, 2010; Nathan, 2002; Oppenheim and Yang, 2005; Yang et al., 2017). However, uncontrolled production of alarmins is harmful or even fatal to the host in some cases. HMGB1 protein acts as a late mediator of lethal systemic inflammation in sepsis (Wang et al., 2004). Therefore, anti-HMGB1 therapeutics have shown to be beneficial in experimental models of sepsis.

S100A8 and S100A9 make up approximately 45% of the cytoplasmic proteins present in neutrophils. Both these proteins are members of the S100 group of proteins. S100A8 and S100A9 are also referred to as MRP8 and MRP14, respectively. Under physiological conditions, massive levels of S100A8 and S100A9 are stored in neutrophils and myeloid-derived dendritic cells, while low levels of S100A8 and S100A9 are expressed constitutively in monocytes (Foell et al., 2004; Wang et al., 2018). S100A8 and S100A9 often form heterodimers (S100A8/A9) (Ometto et al., 2017). The major functions of S100A8/A9 reported so far include the regulation of leukocyte migration and trafficking, the remodeling of cytoskeleton, amplification of inflammation and exertion of anti-microbial activity (Ometto et al., 2017; Wang et al., 2018). After being infected with bacteria, neutrophils, macrophages, and monocytes intensely induce the expression and secretion of S100A8/A9 to modulate inflammatory processes through the induction of inflammatory cytokines. S100A8/9 is an endogenous ligand of toll-like receptor 4 (TLR4) and can trigger multiple inflammatory pathways mediated by TLR4 (Vogl et al., 2007). S100A8 and S100A9 also have antibacterial potential via their ability to bind Zn^2+^ (Foell et al., 2004; Wang et al., 2018). Not much is known about the roles of S100A8/A9 in host defense responses against viruses.

In the present study, we characterized the nature of the early innate immune responses evoked in rhesus macaques and mice during SARS-CoV-2 infection. S100A8 was dramatically upregulated by SARS-CoV-2 and a mouse coronavirus (mouse hepatitis virus, MHV), but not by other viruses. A group of non-canonical neutrophils were also activated during SARS-CoV-2 infection. The abnormal immune responses were mediated by the S100A8/A9-TLR4 pathway. S100A8/A9 specific inhibitor, Paquinimod, significantly reduced the number of neutrophils activated by the coronavirus, inhibited viral replication and rescued lung damage a result of SARS-CoV-2 infection. These results highlight the potential of therapeutically targeting S100A8/A9 for suppressing the uncontrolled inflammation associated with severe cases of COVID-19 and provide new information on an alarmin-mediated pathway for regulating neutrophil function.

## Results

### Induction of S100A8 and activation of neutrophils by SARS-CoV-2

To characterize the early immune responses against coronavirus infection, we infected rhesus macaques with SARS-CoV-2 and analyzed the transcriptome of infected and non-infected animals (Figure 1A). The whole genome wide RNA-seq analysis of the lungs from infected rhesus macaques showed that a number of transcripts were induced or inhibited at day 3 and day 5 after SARS-CoV-2 infection (Supplementary Figure 1A). The induction of primary antiviral factors, like type I IFNs was completely blocked during SARS-CoV-2 infection (Supplementary Figure 1B). Gene ontology (GO) analysis showed that a small group of genes involved in defense responses against viruses was induced (Figure 1B). However, interestingly, a greater number of genes involved in regulating cellular responses to lipopolysaccharide (LPS) were induced by SARS-CoV-2 at day 3 post infection. An analysis of the KEGG pathways also showed that SARS-CoV-2 induced genes were enriched with more from anti-bacterial pathways than those from anti-viral pathways (Supplementary Figure 1C). In line with this, neutrophil chemotaxis associated genes were also upregulated as a result of SARS-CoV-2 infection. This upregulation was less significant at day 3, but more significant at day 5 when compared to the levels of genes related to antiviral responses that were induced as a result of the viral infection (Figure 1B). It is worthy to note that the number of neutrophils increases in COVID-19 patients with severe symptoms (Kuri-Cervantes et al., 2020; Liao et al., 2020; Tan et al., 2020; Wu et al., 2020a). Our results of the preliminary characterization of the early innate response to a SARS-CoV-2 challenge indicate that neutrophils are significantly activated at the very beginning of SARS-CoV-2 infection. To confirm this, we examined the expression of neutrophil markers in the lungs from infected rhesus macaques. All the signature genes, from primary granules to tertiary granules, were significantly induced as a result of SARS-CoV-2 infection (Figure 1C). The markers for monocytes and natural killer cells were slightly upregulated. T cells were unchanged, while interestingly, B cells were significantly down-regulated in the lungs of infected animals (Figure 1D and Supplementary Figure 1D). Taken together, the above data suggested that the SARS-CoV-2 infection provoked a non-canonical antiviral response, or an antibacterial response accompanied by increased neutrophils in the lung at the early stage.

**Fig. 1.**
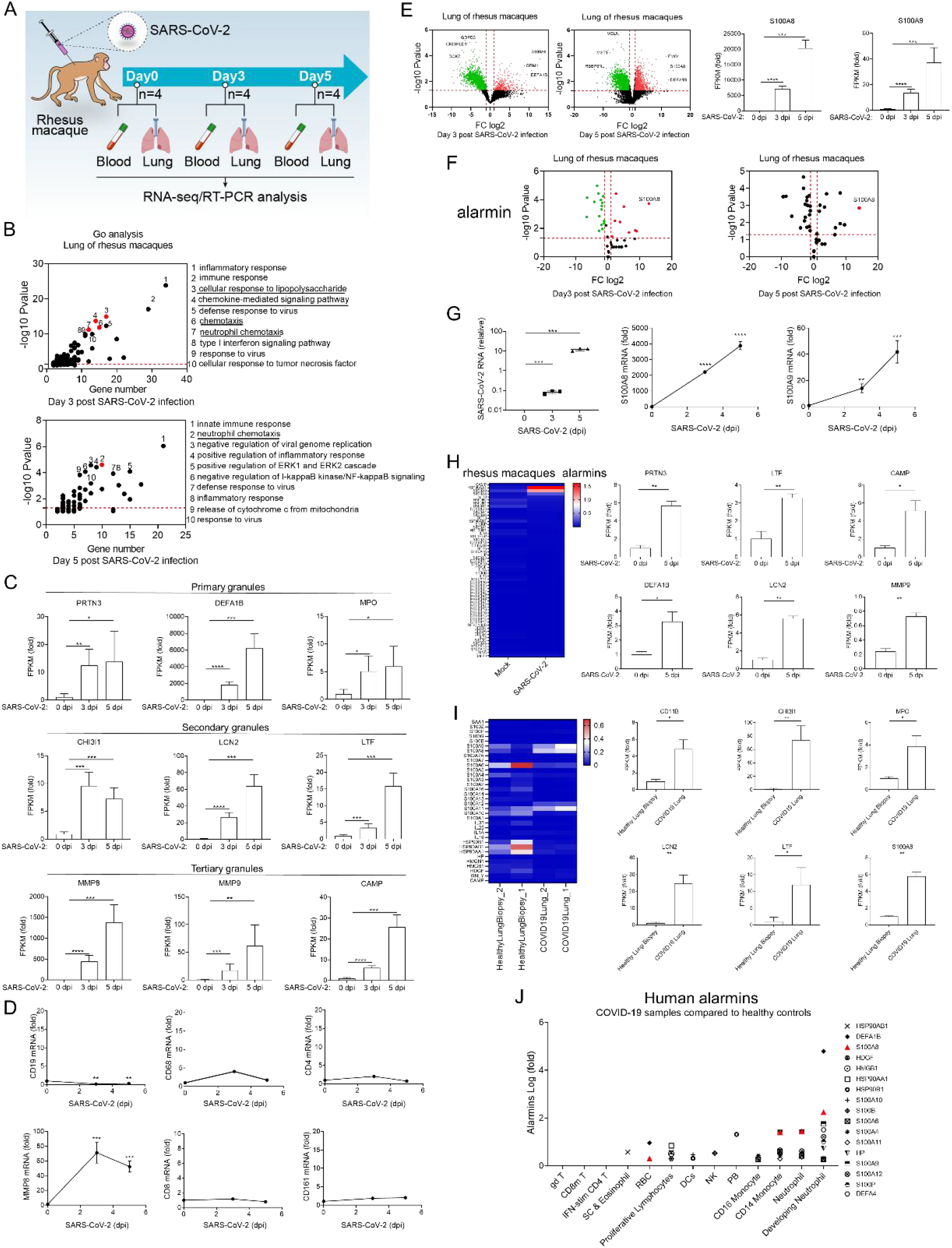
Induction of S100A8 and activation of neutrophils by SARS-CoV-2. **(A)** A flow chart depicting the process of animal experiments with rhesus macaques. **(B)** Go analysis of the differentially expressed genes compared with Mock (*Fold Change (FC) > 4 or < 0*.*25, P value < 0*.*05*). **(C)** The expression of indicated marker genes was analyzed. *n = 4*. **(D)** qRT-PCR analysis for the expression level of *CD19, CD68, CD4, MMP8, CD8*, and *CD161* in the lung of SARS-CoV-2-infected rhesus macaques at 0 dpi, 3 dpi, and 5 dpi. *n = 3*. **(E)** RNA-seq analysis of RNA from lung of rhesus macaques infected intranasally with SARS-CoV-2 at 3 dpi and 5dpi. Differential expressed genes represented by *S100A8* and *S100A9* were analyzed (compared with Mock). **(F)** Analysis of all known alarmins showing that *S100a8* is the most significantly induced one (E). *n = 4*. (**G)** qRT-PCR analysis for viral load and the expression level of *S100*A*8* and S100A9 in the lung of SARS-CoV-2-infected rhesus macaque at 0 dpi, 3 dpi, and 5 dpi. *n = 3*. **(H)** RNA-seq analysis of alarmins from in the blood of rhesus macaques infected intranasally with SARS-CoV-2 at 5dpi. The expression of indicated genes was analyzed (left, relative to *ACTB*; right, fold change to mock). *n = 4*. **(I)** Heatmap depicting the expression levels of alarmins of the samples from health control and post-mortem lung samples from COVID-19-positive patients (relative to *ACTB*, left). The expression of neutrophil marker genes also analyzed (fold change to health control, right). Data from the lung of COVID-19 patients and health control correspond to GEO: GSE147507. **(J)** Analysis of the alarmin expression in peripheral blood from health control and COVID-19 patients. Fold change to health control (log_10_). Data from the peripheral blood of COVID-19 patients and health control correspond to GEO: GSE150728. (*\*P < 0*.*05; **P < 0*.*01; ***P < 0*.*001; ****P < 0*.*0001*).

To explore how an infection by SARS-CoV-2 triggers the activation of antibacterial responses, we examined the differential expression of genes before and after coronavirus infection. The expression of alarmin protein S100A8 was robustly upregulated with 7,000- and 18,000-fold induction at day 3 and day 5 post SARS-CoV-2 infection, respectively (Figure 1E). S100A8 acts as an alarmin through formation of heterodimers with S100A9, then these heterodimers function as danger associated molecular pattern (DAMP) molecules and activate innate immune responses via binding to pattern recognition receptors such as Toll-like receptor 4 (TLR4). In our studies, S100A9 was also induced during the SARS-CoV-2 infection (Figure 1E). The basal expression of S100A9 was much higher, but its induction was not as significant as that of S100A8. In fact, among all the known alarmins, S100A8 was the most significantly induced gene at both day 3 and day 5 post infection (Figure 1F). qRT-PCR analysis showed that the level of S100A8/9 surged along with an increase in the viral load in the lung of infected animals (Figure 1G). Next, we examined the expression of alarmins in the blood samples taken from infected rhesus macaques. S100A8/9 as well as the neutrophil marker genes were also induced by the viral infection (Figure 1H). Based on these results, we investigated if S100A8/A9 were upregulated in COVID-19 patients. Remarkably, both S100A8 and S100A9 were upregulated in post-mortem lung samples from COVID-19-positive patients when compared with biopsied healthy lung tissue from uninfected individuals (Figure 1I). The expression of neutrophil marker genes was also significantly increased in COVID-19-positive patients (Figure 1I). Concomitantly, the mRNA level of S100A8 was significantly higher in peripheral blood from COVID-19-positive patients when compared to healthy subjects (Supplementary Figure 1E). A group of alarmins were induced in different types of blood cells of COVID-19-positive patients. For instance, S100A8 was significantly induced in red blood cells, CD14^+^ monocytes, CD16^+^ monocytes, neutrophils and developing neutrophils (Figure 1J). S100A8/A9 is the ligand of TLR4, which is the primary PRR that recognizes invading gram-negative bacterium and LPS. We thus observed a significant induction of genes that were predominantly involved in responses to LPS and anti-bacterial pathways. S100A8/A9 are also able to induce neutrophil chemotaxis and adhesion, which probably contributed to the increased infiltration of neutrophils.

### Activation of coronavirus-specific neutrophils

To further investigate the nature of immune responses triggered and the roles of S100A8 in these responses, we challenged hACE2 transgenic mice with SARS-CoV-2 and C57BL/6 mice with Influenza A virus (IAV), which also infects the respiratory tract. We performed whole genome wide RNA-seq analysis to characterize the defense responses during viral infection (Figure 2A). Consistent with the results from the rhesus macaque experiments, SARS-CoV-2 infection did not trigger typical antiviral immune responses (Figure 2B and Supplementary Figure 2A). GO analysis showed that SARS-CoV-2 induced genes were enriched in antibacterial humoral response and neutrophil chemotaxis at day 1, day 3 and day 5 post infection. In contrast to this, IAV induced genes were enriched in defense response to virus and cellular response to IFNβ (Figure 2B and Supplementary Figure 2B). KEGG analysis showed that SARS-CoV-2 induced genes were enriched in bacterium related and autoimmune pathways at day 1, day 3 and day 5 post infection. Comparatively, IAV induced genes were enriched within virus related pathways (Figure 2B and Supplementary Figure 2B). We further analyzed the differentially induced genes at different time intervals after SARS-CoV-2 infection. GO analysis showed that while SARS-CoV-2 induced antibacterial responses, neutrophil related and other processes, IAV induced canonical antiviral responses and activated type I IFNs signaling (Figure 2C and supplementary Figure 2C). We then evaluated the expression of IFNs in the lungs during SARS-CoV-2 and IAV infection. Except for IFNκ, all the type I IFNs were significantly induced by IAV infection (Supplementary Figure 2D). Interestingly, SARS-CoV-2 was only able to induce IFNκ but not any other type I IFNs (Supplementary Figure 2D). Both SARS-CoV-2 and IAV were able to induce IFNγ expression. IAV, but not SARS-CoV-2, induced type III IFNs production (Supplementary Figure 2D). qRT-PCR further confirmed that IFNβ and ISG15 induction was impaired during SARS-CoV-2 infection (Figure 2C). The induction of most ISGs was attenuated during SARS-CoV-2 infection compared with IAV infection (Supplementary Figure 2E). However, SARS-CoV-2 but not IAV induced S100A8 and neutrophil marker genes Ly6g expression (Figure 2C). Consistently, other neutrophil marker genes were also induced by SARS-CoV-2 but not by IAV (Figure 2D). Similar to the data from rhesus macaque experiments, compared to other alarmins, S100A8 was robustly induced by SARS-CoV-2 but not by IAV infection in mice (Figure 2E). Thus, SARS-CoV-2 infection specifically induces the transcription of alarmin S100A8. To further confirm this, we infected C57BL/6 mice with other RNA- or DNA-viruses including encephalomyocarditis virus (EMCV), herpes simplex virus 1 (HSV-1) and vesicular stomatitis virus (VSV) and measured the expression of S100A8 in the lung of infected animals. None of these viruses were able to induce the expression of S100A8 (Figure 2F).

**Fig. 2.**
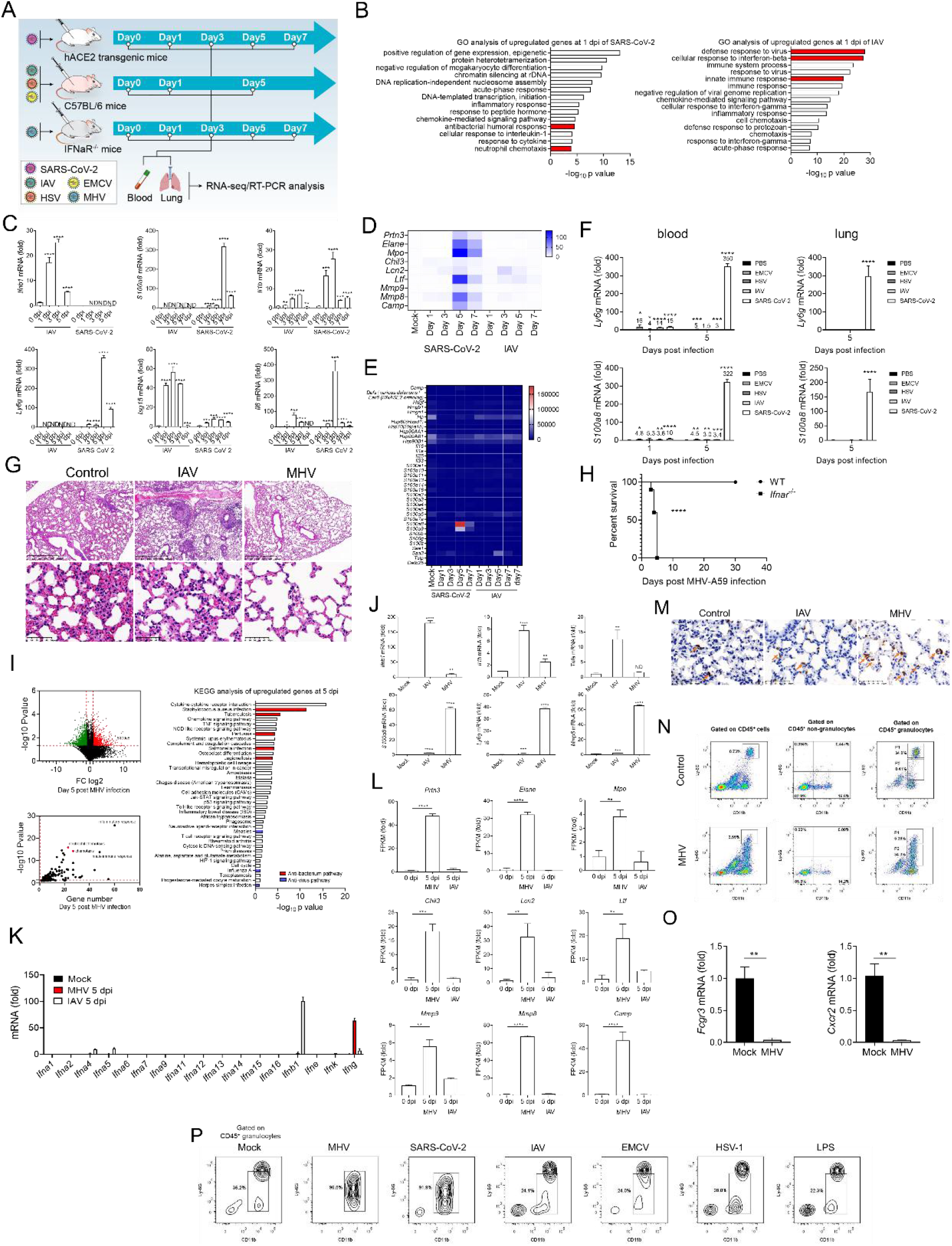
Activation of coronavirus-specific neutrophils. **(A)** A flow chart depicting the process of animal experiments with mice. (**B)** RNA-seq analysis of RNA from lungs of C57BL/6 mice infected with IAV and hACE2 mice infected intranasally with SARS-CoV-2 at 1 dpi. Go analysis was performed with the differentially expressed genes compared with Mock (*FC > 4 or < 0*.*25, P value < 0*.*05*). *n = 3*. (**C)** qRT-PCR analysis for the expression level of *Ifnb1, S100a8, Il6, Isg15, Ly6g* and *Il1b* in the lungs of mice infected intranasally with IAV and SARS-CoV-2 at 0 dpi, 3 dpi, 5 dpi, and 7 dpi. *n = 3*. **(D)&(E)** Heatmap depicting the expression changes of neutrophil marker genes (D) and alarmins (E) in the lungs of mice infected intranasally with IAV and SARS-CoV-2 at 0 dpi, 3 dpi, 5 dpi, and 7 dpi. (**F)** qRT-PCR analysis for the expression level of *S100a8* and *Ly6g* in the blood and lungs of mice infected intranasally with a variety of different viruses (EMCV, HSV-1, IAV, SARS-CoV-2) at 5 dpi, only in the blood and lung of mice infected with SARS-CoV-2 showed a sharp increase in *S100a8* and *Ly6g* expression. (**G)** H&E staining images for the lung of *Ifnar*^-/-^ mice infected intranasally with IAV and MHV at 5 dpi. Higher magnification images of the representative area in lung are shown in the second row. Control means *Ifnar*^-/-^ mice treated intranasally with Vehicle. (**H)** Post-infection survival curves of WT C57BL/6 mice and *Ifnar*^-/-^ mice infected intranasally with MHV. *n = 10/group*. **(I)** RNA-seq analysis of the RNA in the lungs of *Ifnar*^-/-^ mice infected intranasally with MHV at 5 dpi. Go and KEGG analysis was performed with the differentially expressed genes compared with Mock (*FC > 4 or < 0*.*25, P value < 0*.*05*). **(J)** qRT-PCR analysis for the expression level of *Ifnb1, Il1b, Tnfa, S100a8, Ly6g* and *Mmp8* in the lungs of *Ifnar*^-/-^ mice infected intranasally with IAV and MHV at 5 dpi. Mock group means *Ifnar*^-/-^ mice treated intranasally with Vehicle. *n = 3*. **(K)** The expression of *IFNs* in the lungs of *Ifnar*^-/-^ mice infected intranasally with IAV and MHV at 5 dpi was analyzed (Fold change to mock). **(L)** The expression of neutrophil maker genes was analyzed. **(M)** Immunohistochemical staining images for S100A8 in the lung of *Ifnar*^-/-^ mice infected intranasally with IAV and MHV at 5 dpi. The arrows indicate the S100a8-positive cells. Control group means *Ifnar*^-/-^ mice treated intranasally with Vehicle. **(N)** Representative flow cytometry plots defining a special class of neutrophil population (CD45^+^CD11b(high)Ly-6G(mid)) in the lung of *Ifnar*^-/-^ mice infected intranasally with MHV at 5 dpi. Gate P1 and P2 show the difference between special neutrophil population in the lung of *Ifnar*^-/-^ mice infected with MHV and conventional neutrophil population in control group. Control group means *Ifnar*^-/-^ mice treated intranasally with Vehicle. **(O)** qRT-PCR analysis for the expression level of FCGR3and CXCR2 of neutrophil groups (CD11b(high) and Ly-6G(high)) in lung of *Ifnar*^-/-^ mice infected with MHV at 5 dpi. n = 3. **(P)** Flow cytometry plot of lung tissues of hACE2 mice infected intranasally with SARS-CoV-2 and *Ifnar*^-/-^ mice infected intranasally with MHV, IAV, EMCV, HSV-1, and LPS at 5 dpi. Mock group means *Ifnar*^-/-^ mice treated intranasally with Vehicle. Results indicate that the special neutrophil populations appeared only in the mice infected with MHV and SARS-CoV-2. (*\*P < 0*.*05; **P < 0*.*01; ***P < 0*.*001; ****P < 0*.*0001*).

We then investigated if other coronaviruses were able to induce the transcription of S100A8 and activate neutrophils. We infected C57BL/6 mice with mouse hepatitis virus (MHV-A59) intranasally (Figure 2A). Surprisingly, we did not observe any obvious symptoms in infected mice. We then infected IRF3/IRF7 double knockout mice and IFNAR deficient mice with MHV. Similar to the wild type mice, IRF3/IRF7 double knockout mice were able to eliminate the virus rapidly and did not develop severe pneumonia. Interestingly, IFNAR deficient mice showed obvious damage in pulmonary interstitium and alveoli (Figure 2G). All the mice died within 10 days (Figure 2H). We performed whole genome RNA-seq analysis to determine if MHV induced similar immune responses as those induced by SARS-CoV-2 infection. The lungs of MHV infected mice were collected at day 5 post infection and subjected to RNA analysis. Similarly, MHV induced genes were enriched in neutrophil chemotaxis and antibacterial pathways (Figure 2I). S100A8 was also significantly induced by MHV infection (Figure 2J). When compared to IAV, type I IFNs induction was impaired, but neutrophil marker genes were significantly induced by MHV infection (Figure 2K and 2L). Histological staining of samples taken from lungs showed more S100A8 positive cells infiltration after MHV infection (Figure 2M).

Since the expression of neutrophil marker genes was elevated in the lung after coronavirus infection, we examined if more neutrophils were infiltrated into lung during infection. Surprisingly, the number of Ly-6G (high) and CD11b (high) neutrophils decreased in the lung after MHV infection (Figure 2N). Interestingly, a group of CD11b (high) Ly-6G (mid) cells showed up after MHV infection. Further analysis revealed that these cells were CD45^+^ granulocytes. A recent single cell analysis study had reported that a group of developing neutrophils was detected in the peripheral blood of COVID-19 patients with decreased expression of canonical neutrophil markers CXCR2 and FCGR3B (Wilk et al., 2020). We purified these cells by using flow cytometry sorting. qRT-PCR analysis showed decreased expression of CXCR2 and FCGR3 (Figure 2O). To explore if other types of viruses or stimuli are able to induce these cells, we challenged C57BL/6 mice with IAV, EMCV, VSV, HSV-1 and LPS, hACE2 transgenic mice with SARS-CoV-2 through nasal inoculation. Interestingly, only SARS-CoV-2 and MHV induced this group of neutrophils (Figure 2P). Taken together, these results suggest that coronavirus infection induces S100A8 and a group of noncanonical neutrophils specifically.

### Paquinimod suppresses coronavirus specific neutrophils and viral infection

S100A8/A9 functions as a TLR4 ligand that amplifies inflammation by triggering the activation of innate immune signaling, including neutrophil chemotaxis. We thus reasoned that the abnormal upregulation of S100A8/A9 probably resulted in activation of coronavirus specific neutrophils and dysregulation of immune responses. Quinoline-3-carboxamide Paquinimod (also known as ABR-215757), is an inhibitor of S100A9 which can prevent the binding of S100A9 to TLR-4 (Bjork et al., 2009; Schelbergen et al., 2015), suggesting that it can be used to block the function of S100A8/A9 and mitigate the antibacterial immune responses elicited during SARS-CoV-2 infection. To test this, we treated the SARS-COV-2 infected mice with S100A8/A9 inhibitor Paquinimod (Figure 3A). Expectedly, Paquinimod was able to rescue the weight loss of infected mice and inhibited SARS-CoV-2 viral replication significantly (Figure 3B and 3C). Consistently, the expressions of neutrophil markers (Ly6G, MMP8, LTF), IL-6 and S100A8 were significantly reduced after Paquinimod treatment (Figure 3D). More strikingly, Paquinimod completely rescued the death of MHV infected, but not IAV infected IFNAR deficient mice and significantly inhibited MHV replication (Figure 3E, 3F and Supplementary Figure 3A). The expression of neutrophil marker genes in lung and blood from infected mice was also inhibited by Paquinimod (Figure 3G, 3H and Supplementary Figure 3B). The damages to pulmonary interstitium and alveoli were alleviated, and infiltration of S100A8^+^ cells was also reduced by Paquinimod (Figure 3I and 3J).

**Fig. 3.**
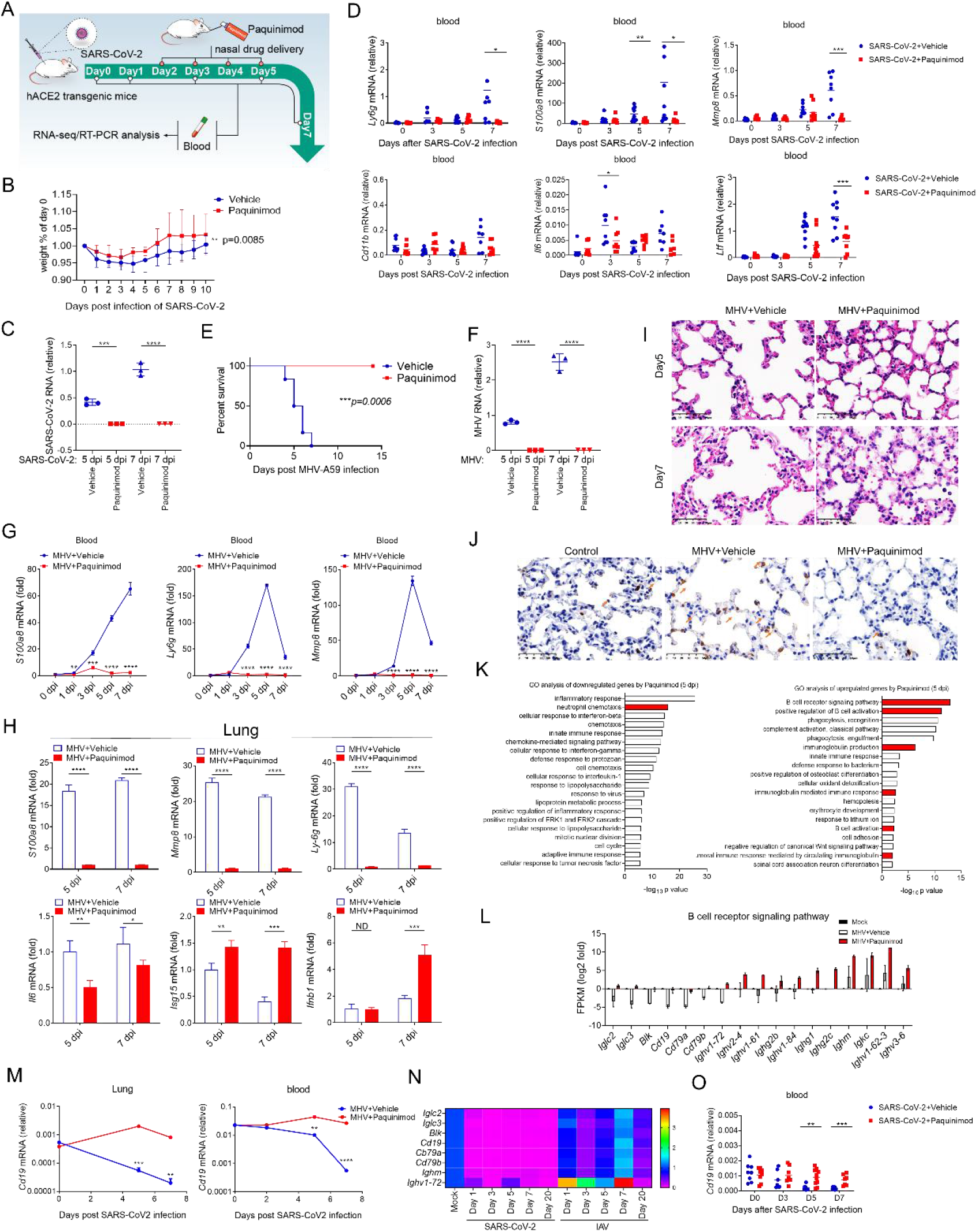
Paquinimod suppresses coronavirus specific neutrophils and viral infection. **(A)** A flow chart depicting the process of animal experiments with mice. **(B)** Percentages of body weight change of hACE2 mice infected intranasally with SARS-CoV-2 between Vehicle treated group and Paquinimod treated group. *n=6/group*. **(C)** qRT-PCR analysis for viral load in the lungs of hACE2 mice infected intranasally with SARS-CoV-2 between Vehicle treated group and Paquinimod treated group at 5 dpi and 7 dpi. *n = 3*. **(D)** qRT-PCR analysis for the expression level of *Ly6g, S100a8, Mmp8, Cd11b, Il6*, and *Ltf* in the peripheral blood from hACE2 mice infected intranasally with SARS-CoV2 between Vehicle treated group and Paquinimod treated group at 0 dpi, 1 dpi, 3 dpi, 5 dpi, and 7 dpi. *n = 3*. **(E)** Post-infection survival curves of *Ifnar*^*-/-*^ mice infected intranasally with MHV between Vehicle treated group and Paquinimod treated group. *n=6/group*. **(F)** qRT-PCR analysis for viral load in the lungs of mice infected intranasally with MHV at 5 dpi and 7 dpi between Vehicle treated group and Paquinimod treated group. *n = 3*. **(G)** qRT-PCR analysis for the expression level of *S100a8, Ly6g* and *Mmp8* in the blood of *Ifnar*^-/-^ mice infected intranasally with MHV at 0 dpi, 1 dpi, 3 dpi, 5 dpi and 7 dpi between Vehicle treated group and Paquinimod treated group. *n = 3*. **(H)** qRT-PCR analysis for the expression level of *Ifnb1, Il1b, Il6, S100a8, Ly6g* and *Mmp8* in the lungs of *Ifnar*^-/-^ mice infected intranasally with MHV at 5 dpi and 7 dpi between Vehicle treated group and Paquinimod treated group. *n = 3*. (**I)** H&E staining images for the lungs of *Ifnar*^-/-^ mice infected intranasally with MHV between Vehicle treated group and Paquinimod treated group at 5 dpi and 7 dpi. **(J)** Immunohistochemical staining images for S100A8 in the lung of *Ifnar*^-/-^ mice between Vehicle treated group and Paquinimod treated group at 5 dpi. The arrows indicate the S100a8-positive cells. **(K)&(L)** RNA-seq analysis of RNA from lung of *Ifnar*^-/-^ mice infected intranasally with MHV between Vehicle treated group and Paquinimod treated group at 5 dpi. Go analysis was performed with the differentially expressed genes (K) (*FC > 4 or < 0*.*25, P value < 0*.*05*), and the expression level of B cell receptor related genes was analyzed (L). **(M)** qRT-PCR analysis for the expression level of *Cd19* in the peripheral blood and lung of *Ifnar*^-/-^ mice infected intranasally with MHV between Vehicle treated group and Paquinimod treated group at 0 dpi, 5 dpi, and 7 dpi. *n = 3*. **(N)** Heatmap depicting the expression level of indicated genes in the lung of mice infected intranasally with SARS-CoV-2 and IAV. **(O)** qRT-PCR analysis for the expression level of *Cd19* in the peripheral blood of hACE2 mice infected intranasally with SARS-CoV2 between Vehicle treated group and Paquinimod treated group at 0 dpi, 1 dpi, 3 dpi, 5 dpi, and 7 dpi. *n = 3*. (*\*P < 0*.*05; **P < 0*.*01; ***P < 0*.*001; ****P < 0*.*0001*).

To gain further insights into the modulation of immune responses after Paquinimod treatment, we performed genome wide RNA-seq analysis using the RNA from MHV infected IFNAR deficient mice. As expected, pathways related to neutrophil chemotaxis and antibacterial responses were primarily downregulated by Paquinimod (Figure 3K and Supplementary Figure 3C). Interestingly, Paquinimod upregulated genes that were intensively enriched in positive regulation of B cell activation and B cell receptor signaling pathway (Figure 3K). The transcription of a group of B cell related genes was decreased during MHV infection (Figure 3L), which was consistent with results in rhesus macaque experiments (Supplementary Figure 1D). The expression of these B cell related genes was rescued or induced by Paquinimod during MHV infection, which was confirmed by qRT-PCR analysis (Figure 3M). In line with this, SARS-CoV-2 but not IAV infection, suppressed the expression of these B cell related genes (Figure 3N and Supplementary Figure 3D), which were also rescued by Paquinimod (Figure 3O).

### S100A8 activates aberrant neutrophils via TLR4

We demonstrated that Paquinimod inhibited the expression of neutrophil maker genes in lung and blood. To determine if Paquinimod could reduce the number of these abnormal cells, we performed flowcytometry to analyze the cell population during coronavirus infection. Coronavirus-specific neutrophils were significantly reduced in blood and lung after treatment with Paquinimod (Figure 4A). It is believed that S100A8/A9 can activate TLR4 or receptor for advanced glycation end products (RAGE) pathways. Interestingly, TLR4 inhibitor (Resatorvid) but not RAGE inhibitor (Azeliragon) was able to reduce the number of these neutrophils suggesting that S100A8/A9 activated these neutrophils via TLR4 pathway (Figure 4A). Moreover, MHV infection also induced these neutrophils in liver and bone marrow, which could be suppressed by TLR4 inhibitor (Figure 4B). Consistently, Resatorvid was able to inhibit viral replication in lung and blood of the infected mice (Figure 4C). Moreover, both Paquinimod and Resatorvid suppressed the activation of coronavirus related neutrophils in lung during SARS-CoV-2 infection (Figure 4D). To confirm the role of TLR4 in activating S100A8 related signaling, we examined S100A8 induction upon LPS treatment. LPS induced S100A8 expression through adaptor protein MyD88 (Supplementary Figure 4A). We then treated mouse macrophages with the serum from SARS-CoV-2 infected mice with or without Paquinimod treatment. The serum from infected mice was able to induce the expression of S100A8 and CXCL2 but not IL-1β or IL-6 in a MyD88 dependent manner (Figure 4E and Supplementary Figure 4B). S100A8 further activated TLR4 signaling to form a positive loop. We thus observed that Paquinimod was able to inhibit the induction of S100A8 (Figure 3D and 3G-H). CXCL2 is the primary chemokine that recruits neutrophils. Consistent with the *in vivo* results (Figure 4A), Azeliragon could not inhibit induction of S100A8 or CXCL2 by the serum (Figure 4E). S100A8 therefore exerted its pathogenic function through activating TLR4 signaling during coronavirus infections.

**Fig. 4.**
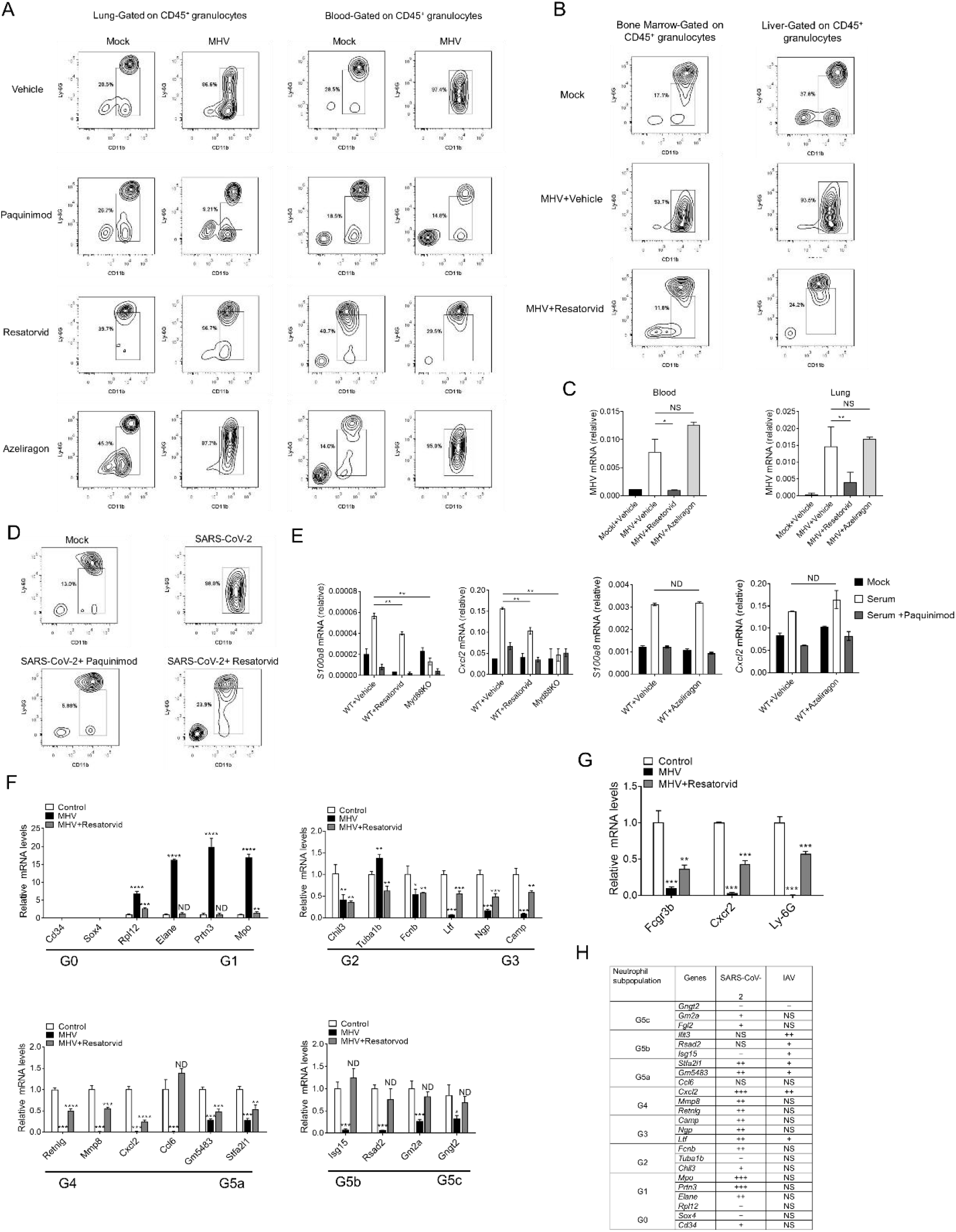
S100A8 mediates activation of aberrant neutrophils via TLR4. **(A)** Flow cytometry detecting coronavirus specific neutrophils in the lung and blood of *Ifnar*^-/-^ mice infected with MHV among Vehicle treated group, Paquinimod treated group, Resatorvid treated group, and Azeliragon treated group at 5 dpi. **(B)** Flow cytometry detecting coronavirus specific neutrophils in bone marrow and liver of *Ifnar*^-/-^ mice infected with MHV between control group and Resatorvid treated group at 7 dpi. Mock group means *Ifnar*^-/-^ mice treated intraperitoneal with Vehicle. **(C)** qRT-PCR analysis for viral load in the lungs of mice infected intranasally with MHV at 5 dpi between Vehicle treated group and Resatorvid treated group. *n = 3*. **(D)** Flow cytometry detecting coronavirus specific neutrophils in the lungs of hACE2 mice infected with SARS-CoV-2 between control group, Paquinimod and Resatorvid treated group at 5 dpi. **(E)** qRT-PCR analysis for the expression level of *Cxcl2* and *S100a8* in the Raw 264.7 cells treated with indicated serum among WT control group, WT Resatorvid treated group, WT Azeliragon treated group and *Myd88*^-/-^ group. n = 3. **(F)&(G)** qRT-PCR analysis for the expression level of several related genes of neutrophil groups in bone marrow of *Ifnar*^-/-^ mice infected with MHV between control group and Resatorvid treated group at 5 dpi. n = 3. **(H)** RNA-seq analysis of the level of several related genes of neutrophil groups from the lungs of C57BL/6 mice infected with IAV and hACE2 mice infected with SARS-CoV-2 at 5 dpi. (*\*P < 0*.*05; **P < 0*.*01; ***P < 0*.*001; ****P < 0*.*0001*).

A recent single cell sequencing data clarified the heterogeneity of neutrophils and identified 8 subpopulations (Xie et al., 2020). We collected the CD11^+^ and Ly-6G^+^ cells from bone marrow (Figure 4B) and examined the expression of each marker gene during MHV infection with or without TLR4 inhibitor. Compared to noninfected mice, MHV induced expression of all 3 G1 maker genes and 1 maker gene from G0 to G1, but inhibited expression of other maker genes (Figure 4F). Interestingly, Resatorvid was able to reverse the expression of marker genes back to the normal levels. The expression of canonical neutrophil markers CXCR2, LY6G and FCGR3 were decreased after MHV infection (Figure 4G). We further analyzed the RNA-seq data of lung samples taken from the mice infected with SARS-CoV-2 and IAV. G5B population, the matured neutrophil, was activated during IAV infection (Figure 4H and Supplementary Figure 4C). Whereas SARS-CoV-2, similar to the results from bone marrow experiments (Figure 4F), activated G1 subpopulation significantly. In addition, SARS-CoV-2 also induced other premature neutrophil markers from G2 to G4 subpopulation in lung (Figure 4H and Supplementary Figure 4C). Interestingly, Paquinimod was able to turn off the abnormal upregulation of these genes (Supplementary Figure 4D). Together, these results suggested that coronavirus infection expanded the population of premature neutrophils in S100A8/9 and TLR4 dependent manner, and caused dysregulation of the immune system.

## Discussion

Innate immunity, the first line of host defense, employs pattern-recognition receptors (PRRs) for recognizing pathogen-associated molecular patterns (PAMPs) of invading pathogens and galvanizing the host defense machinery. The endogenous danger-associated molecular patterns (DAMPs) are also able to trigger the activation of innate immune signaling. Alarmins are a panel of proteins or peptides that can function as DAMPs to activate various immune pathways (Bianchi, 2007; Yang et al., 2017). The fine tuning of the transcription of alarmins is critical for maintaining immune homoeostasis. Over or sustained expression of alarmins could result in uncontrolled inflammation (Chan et al., 2012; Cher et al., 2018; Kang et al., 2014; Patel, 2018). Such imbalanced immune responses and cytokine storm contribute to the development of severe acute respiratory distress syndrome (ARDS). Here we demonstrated that SARS-CoV-2 and MHV, but not any other tested viruses, induced a robust and sustained transcription of the alarmin S100A8 in animal models (Figure 1 D-G and Figure 2). In line with this, the substantial upregulation of S100A8 was also observed in the lung and peripheral blood of infected macaques and COVID-19 patients (Figure 1H and 1I). Although we did observe that the SARS-CoV-2 and the other types of viruses could induce S100A8 expression slightly in blood at day 1 post infection (Figure 2C), however, the induction of S100A8 by viruses other than SARS-CoV-2 was either unchanged or reduced at day 5 post infection. S100A8 expression increased dramatically by 300 folds at day 5 post infection (Figure 2C). Similar phenotypes were observed during MHV infection (Figure 2F). Thus, uncontrolled or sustained induction of S100A8 could be a common feature of coronavirus infections. However, the mechanisms by which coronaviruses provoke the induction of S100A8 is unknown.

In addition to studying the innate immune responses elicited by SARS-CoV-2 infected hACE2 transgenic mice and rhesus macaques, we also infected IFNAR deficient mice with mouse coronavirus and characterized the resulting immune responses. The MHV infected mice exhibited a characteristic immune signature comprising of S100A8 induction, initiation of antibacterial like responses and activation of coronavirus related neutrophils. These features of the immune response were similar to those observed in SARS-CoV-2 infected mice and rhesus macaques (Figure 2 and Figure 4A and 4B). Untreated MHV infected mice developed more severe symptoms and died within 10 days. Mice administered with S100A8 inhibitor, Paquinimod, were rescued from lung damage and death against MHV infections. Since these results were similar to those obtained during studies with SARS-CoV-2, it seems like a shared mechanism that directs the pathogenesis of pneumonia during SARS-CoV-2 and MHV infections in animal models. Thus, IFNAR deficient mice and MHV through its ability to evoke similar immune responses as SARS-CoV-2, could serve as useful models for investigating ARDS associated with SARS-CoV-2 infection. Although we inoculated wild type, MAVS knockout, STING knockout, ELF4 knockout and IRF3/IRF7 double knockout mice with MHV intranasally, none of these mice developed obvious ARDS related symptoms. IRF3 and IRF7 are key transcription factor of type I IFNs. Type I IFNs activate interferon stimulated genes by binding IFNAR. Since the IRF3/IRF7 double knockout phenotype was not susceptible to MHV infection, it seems like IFNAR can exert its function in a type I IFNs independent way. Previous studies (Hadjadj et al., 2020; Zhou et al., 2020) and our above data showed that induction of type I IFNs is completely blocked during SARS-CoV-2 and MHV infections. This further suggests that IFNAR is involved in defense against coronavirus with an unknown mechanism.

There is consensus that the neutrophil number is significantly increased in COVID-19 patients with severe symptoms (Kuri-Cervantes et al., 2020; Liao et al., 2020; Tan et al., 2020; Wu et al., 2020a). Moreover, a recent study identified that a group of premature neutrophils in COVID-19 patients that do not express canonical neutrophil markers like CXCR2 and FCGR3B. Bacterial coinfection is common in viral pneumonia, especially, in critically ill patients (Bost et al., 2020). Neutrophils can be activated during bacterial infection and are critical for killing invading bacteria (Deng et al., 2013; Li et al., 2002). The increased neutrophil number and developing neutrophils in patients could be attributed to coinfection of bacterium. However, results of our studies showed that the premature neutrophils were induced by coronavirus at the early stage of infection in animal models. We compared these cells with 8 neutrophil subpopulations that were proposed in a recent study. Coronavirus induced neutrophils in bone marrow and lung belong to the G1 population predominately (Figure 4E and Supplementary Figure 4B). Furthermore, the populations of neutrophils in lungs are more diverse than those in bone marrow. Compared to IAV induced neutrophils, coronavirus preferentially induces the expression of premature marker genes (Supplementary Figure 4B). However, the exact mechanism by which coronavirus induces these abnormal neutrophils and the key transcription factors that direct development of these cells await to be further investigated.

In summary, we have demonstrated that alarmin S100A8 was specifically upregulated by SARS-CoV-2 infection. S100A8/A9 amplified inflammatory responses by activating TLR4 signaling pathway. A group of coronavirus-specific premature neutrophils were activated during infection. The inhibitors of S100A8/A9-TLR4 axis were able to mitigate abnormal inflammation and inhibit viral replication. These results uncover the role of alarmins and neutrophils in the pathogenesis of SASR-CoV-2 infection and provide new therapeutic targets for the treatment of COVID-19.

## Acknowledgments

This work was supported by the National Natural Science Foundation of China (31570891; 31872736), the National Key Research and Development Program of China (2016YFA0500302; 2020YFA0707800), the National Key Research and Development Program (2020YFA0707500) and the Strategic Priority Research Program (XDB29010000). Xiangxi Wang was supported by Ten Thousand Talent Program and the NSFS Innovative Research Group (81921005). We thank National Mega projects of China for Major Infectious Diseases (2017ZX10304402), CAMS initiative for Innovative Medicine of China (2016-12M-2-006) and The National Natural Science Foundation of China (82041008) for the support on the animal model study.

## Author Contributions

F.Y., X.W., C.Q. and Q.G. conceived the study and analyzed the data. Q.G., Y.Z. and J.L. performed most experiments and analyzed the data. F.Y. and X.G. analyzed the RNA-seq data. Z.Z., L.C. and Y.L. helped with the mice experiments. Y.L., X.W, X. Wei, L. Chen. provided support on literature search. F.Y. wrote the paper. F.Y., X.W. and C.Q. revised the paper.

## STAR Methods

### Facility and ethics statements

All experiments with live SARS-CoV-2 viruses were carried out in the enhanced biosafety level 3 (P3+) facilities in the Institute of Laboratory Animal Science, Chinese Academy of Medical Sciences (CAMS) approved by the National Health Commission of the People’s Republic of China. All animals care and use were in accordance with the Guide for the Care and Use of Laboratory Animals of the Chinese Association for Laboratory Animal Science. All procedures of animal handling were approved by the Animal Care Committee of Peking University Health Science Center.

### Animal experiments

All animals were kept and bred in specific pathogen-free conditions. Wild-type (WT) mice were purchased from Department of Laboratory Animal Science of Peking University Health Science Center, Beijing. Male and female hACE2 mice on a C57BL/6J background were obtained from the Institute of Laboratory Animal Science, Peking Union Medical College. *Elf4*^-/-^ mice, *Ifnar*^-/-^ mice and *Mavs*^-/-^ mice have been described previously. *Irf3/7* double knockout mice on a C57BL/6J background was a gift from Pro. Zhengfan Jiang.

Before being inoculated intranasally with reagents and viruses, hACE2 mice were intraperitoneally anaesthetized by 2.5% avertin with 0.02 mL/g body weight; WT C57BL/6J mice, interferon-α receptor gene knockout mice (*Ifnar*^-/-^), *Elf4*^-/-^ mice, *Ifnar*^-/-^ mice and *Mavs*^-/-^ mice were anaesthetized by isoflurane; rhesus macaques were anaesthetized with 10 mg/kg ketamine hydrochloride. The health status and body weight of all mice were observed and recorded daily. Mice were euthanized at 0, 1, 3, 5 and 7 dpi to collect different tissues and examine virus replication and histopathological changes.

### Viruses

VSV (Vesicular Stomatitis Virus, Indiana strain) was a gift from J. Rose (Yale University), IAV (Influenza A Virus, PR8) was a gift from Feng Qiang (Fudan University), and HSV-1 (Herpes simplex virus 1) was from A. Iwasaki (Yale University). EMCV (Encephalomyocarditis virus, VR-129B) was purchased from American Type Culture Collection (ATCC). MHV-A59 (mouse hepatitis virus A-59) has been described previously (Yang et al., 2014). The SARS-CoV-2 which has been used in this paper is named strain HB-01. The complete genome for this SARS-CoV-2 had been put in to GISAID (BetaCoV/Wuhan/IVDC-HB-01/2020|EPI_ISL_402119), and has been described previously (Yu et al., 2020).

VSV, EMCV, HSV-1 were propagated in Vero cells followed by 3 cycles of freezing and thawing, then the large debrises were spun down and the supernatants were used as a stock solution. The titer of the viruses was determined by plaque assay in Vero cells. For mice infection assay, WT mice were inoculated intranasally with VSV (10^7^ PFU), EMCV (10^7^ PFU), HSV-1 (10^6^ PFU) after anesthesia. PFU: plaque forming unit.

MHV-A59 were propagated in 17CL-1 cells followed by 3 cycles of freezing and thawing, the large debrises were spun down and the supernatants were used as a stock solution. The titer of the viruses was determined by plaque assay in 17CL-1 cells. For mice infection assay, *Ifnar*^-/-^ mice were inoculated intranasally with MHV-A59 (10^5^ PFU) after anesthesia or *Ifnar*^-/-^ mice were inoculated intraperitoneally with MHV-A59 (10^6^ PFU).

IAV was propagated in 10-day-old specific-pathogen-free embryonic chicken eggs. The allantoic fluid was collected and titrated to determine the 50% tissue culture infection dose (TCID50) in A549 cells and the median lethal dose (LD50) in mice following the Reed-Muench method. For mice infection assay, WT mice were inoculated intranasally with IAV (10^5^ TCID50) after anesthesia.

### Paquinimod rescue assay

For the Paquinimod rescue assay, WT mice, hACE2 mice, and *Ifnar*^-/-^ mice were inoculated intranasally with IAV, SARS-CoV-2, and MHV-A59 respectively after anesthesia. The mice were given intranasally 12.5 μg/day of Paquinimod (TargetMol; Catalog No. T7310) starting on 2 dpi. The control group mice were given intranasally 50 μL of PBS. Stock solutions of 100 mg/mL Paquinimod were prepared with DMSO in advance. The health status and body weight of all mice were observed and recorded daily. Mice were euthanized at 0,1, 3, 5 and 7 dpi to collect different tissues and examine virus replication and histopathological changes.

### Resatorvid rescue assay/Azeliragon rescue assay

*Ifnar*^-/-^ mice were inoculated intraperitoneally with MHV-A59. For the Resatorvid rescue assay, the mice were given intraperitoneally 50 μg/day of Resatorvid (MCE; Synonyms: TAK-242; CLI-095) starting on 2 dpi. Stock solutions of 10 mM Resatorvid were prepared with DMSO in advance. Before inoculating, stock solutions of Resatorvid in DMSO was diluted by corn oil. For the Azeliragon rescue assay, the mice were given intraperitoneally 100 μg/day of Azeliragon (TargetMol; Catalog No. T2507) starting on 2 dpi. Stock solutions of 10 mM Azeliragon were prepared with DMSO in advance. Before inoculating, stock solutions of Azeliragon in DMSO was diluted by corn oil. The control group mice were given intraperitoneally 200 μL of corn oil solution which is contained 20 μL DMSO. The health status and body weight of all mice were observed and recorded daily. Mice were euthanized at 0,1, 3, 5 and 7 dpi to collect different tissues and examine virus replication and histopathological changes.

### Cell culture

Raw 274.7 cells, 17CL-1 cells, MDCK cells and Vero cells were kept in our lab. 17CL-1 cells, MDCK cells and Vero cells were cultured in DMEM medium (Gibco) supplemented with 10% FBS (PAN), 100 U/mL Penicillin-Streptomycin. Raw 274.7 cells were cultured in 1640 medium (Gibco) supplemented with 10% FBS, 100 U/mL Penicillin-Streptomycin. Cells were negative for mycoplasma.

### Serum co-culture assay

For Serum co-culture assay, 100 μL peripheral blood was collected from each group of *Ifnar*^-/-^ mice on day 5 during Paquinimod rescue assay. After blood coagulation at room temperature, samples were spun down and the supernatants were used as a stock serum.

Raw 264.7 cells were seeded on 6-well plates with 10^6^ cells/mL. After cell adherence, LPS (100 ng/mL) and mice serum which has been collected above (2 μL/mL) were added. After 12 hours co-culture, cells were harvested and lysed by TRNzol reagent for RNA extraction.

### RNA sequencing (RNA-seq) and analysis

Whole RNA of cells with specific treatment were purified using RNeasy Mini Kit (Qiagen NO. 74104). The transcriptome library for sequencing was generated using VAHTSTM mRNA-seq v2 Library Prep Kit for Illumina® (Vazyme Biotech Co.,Ltd, Nanjing, China) following the manufacturer’s recommendations. After clustering, the libraries were sequenced on Illumina Hiseq X Ten platform using (2×150bp) paired-end module. The raw images were transformed into raw reads by base calling using CASAVA (http://www.illumina.com/support/documentation.ilmn). Then, raw reads in a fastq format were first processed using in-house perl scripts. Clean reads were obtained by removing reads with adapters, reads in which unknown bases were more than 5% and low quality reads (the percentage of low quality bases was over 50% in a read, we defined the low quality base to be the base whose sequencing quality was no more than 10). At the same time, Q20, Q30, GC content of the clean data were calculated (Vazyme Biotech Co.,Ltd, Nanjing, China). The original data of the RNA-seq was uploaded to the GEO DataSets.

### Quantitative RT-PCR (qRT-PCR) analysis

Total RNA was isolated from the tissues by TRNzol reagent (DP424, Beijing TIANGEN Biotech, China). Then, cDNA was prepared using HiScript III 1st Strand cDNA Synthesis Kit (R312-02, Nanjing Vazyme Biotech, China). qRT-PCR was performed using the Applied Biosystems 7500 Real-Time PCR Systems (Thermo Fisher Scientific, USA) with SYBR qPCR Master mix (Q331-02, Nanjing Vazyme Biotech, China). The data of qRT-PCR were analyzed by the Livak method (2^−ΔΔCt^). Ribosomal protein L19 (RPL19) was used as a reference gene for mice, and GAPDH for macaques. qRT-PCR primers are displayed in supplementary materials Table S1.

### Histology and Immunohistochemical staining

The lungs were quickly placed in cold saline solution and rinsed after they were collected. Then, lungs were fixed in 4% PFA, dehydrated and embedded in paraffin prior to sectioning at 4 μm and sections were stained with hematoxylin and eosin (H&E).

For immunohistochemical staining, the lung paraffin sections were dewaxed and rehydrated through xylene and an alcohol gradient. Antigen retrieval was performed by heating the sections to 100 °C for 4 min in 0.01 M citrate buffer (pH 6.0) and repeated 4 times. The operations were performed according to the instructions of the two-step detection kit (PV-9001, Beijing ZSGB Biotechnology, China). The samples were treated by endogenous peroxidase blockers for 10 min at room temperature followed by incubation with primary antibodies S100a8 (1:200, 47310T, Cell Signaling Technology) at 37°C for 1 h, then after washed with PBS, sections were incubated with reaction enhancer for 20 min at room temperature and secondary antibodies at 37°C for 20 min, and finally sections were visualized by 3,30-diaminobenzidine tetrahydrochloride (DAB) and counterstained with haematoxylin.

### Tissue preparation and data analysis for flow cytometry

The lung tissues, peripheral blood and bone marrow were collected from the mice. The lungs were first ground with 200 mesh copper sieve, and then transferred to DMEM containing 10% FBS, 0.5 mg/mL Collagenase D (11088858001, Roche, Switzerland) and 0.1 mg/mL DNase I (07469, STEMCELL Technologies, Canada) for a 20 min digestion at 37°C to obtain single-cell suspensions. Livers were harvested and homogenized into single-cell suspensions using 200 mesh copper sieve. Bone marrow were flushed out of the femurs using a 23–gauge needle in PBS containing 2mM EDTA and 2% fetal bovine serum (FBS) and dispersed into single cells through a pipette. Single-cell samples were treated by red blood cell lysis buffer (R1010, Beijing Solarbio Science & Technology, China) for 2 min at room temperature and passed through a 70–μm nylon mesh sieve before staining. Peripheral blood was treated with red blood cell lysis buffer to remove red blood cells.

After blocking non-specific Fc-mediated interactions with CD16/CD32 antibodies (14-0161-82, eBioscience, USA), single-cell suspensions were stained with fluorophore-conjugated anti-mouse antibodies at 4°C for 30min. After washing the samples, flow cytometry acquisition was performed on a BD LSRFortessa. Sorting were performed using a BD AriaIII (BD). All antibodies were purchased from eBioscience: CD45-PE (12-0451-81), Ly-6G-APC (17-9668-80), CD11b-FITC (11-0112-81).

### Statistical analysis

All analyses were repeated at least three times, and a representative experimental result was presented. Two-tailed unpaired Student’s t test was used for statistical analysis to determine significant differences when a pair of conditions was compared. Asterisks denote statistical significance (*\*P < 0*.*05; **P < 0*.*01; *** P < 0*.*001*). The data are reported as the mean ± S.D.

## Supplemental Information

**Figure S1.**
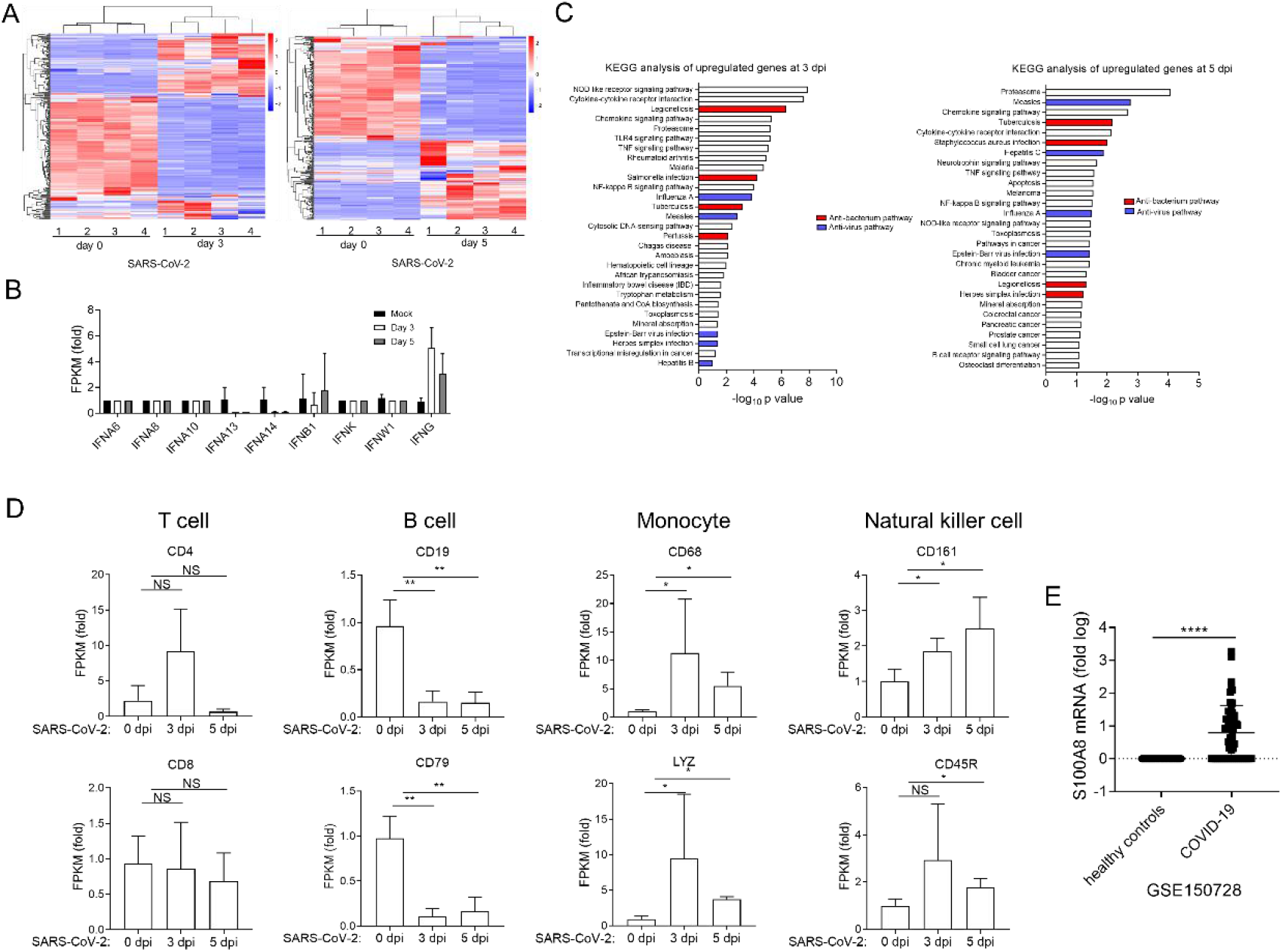
**(A)-(D)** RNA-seq analysis of RNA from lungs of rhesus macaques infected intranasally with SARS-CoV-2 at 3 dpi and 5dpi. Heatmap depicting the differentially expressed genes (A). The expression of IFNs was analyzed (B). KEGG analysis was performed with the differentially expressed genes compared with Mock (C) (*FC > 4 or < 0*.*25, P value < 0*.*0*5). The expression of indicated marker genes was analyzed (D). *n = 4*. **(E)** Analysis of *S100A8* expression in peripheral blood from health control and COVID-19 patients. Fold change to health control (log_10_). Data from the peripheral blood of COVID-19 patients and health control correspond to GEO: GSE150728. (*\*P < 0*.*05; **P < 0*.*01; ***P < 0*.*001; ****P < 0*.*0001*).

**Figure S2.**
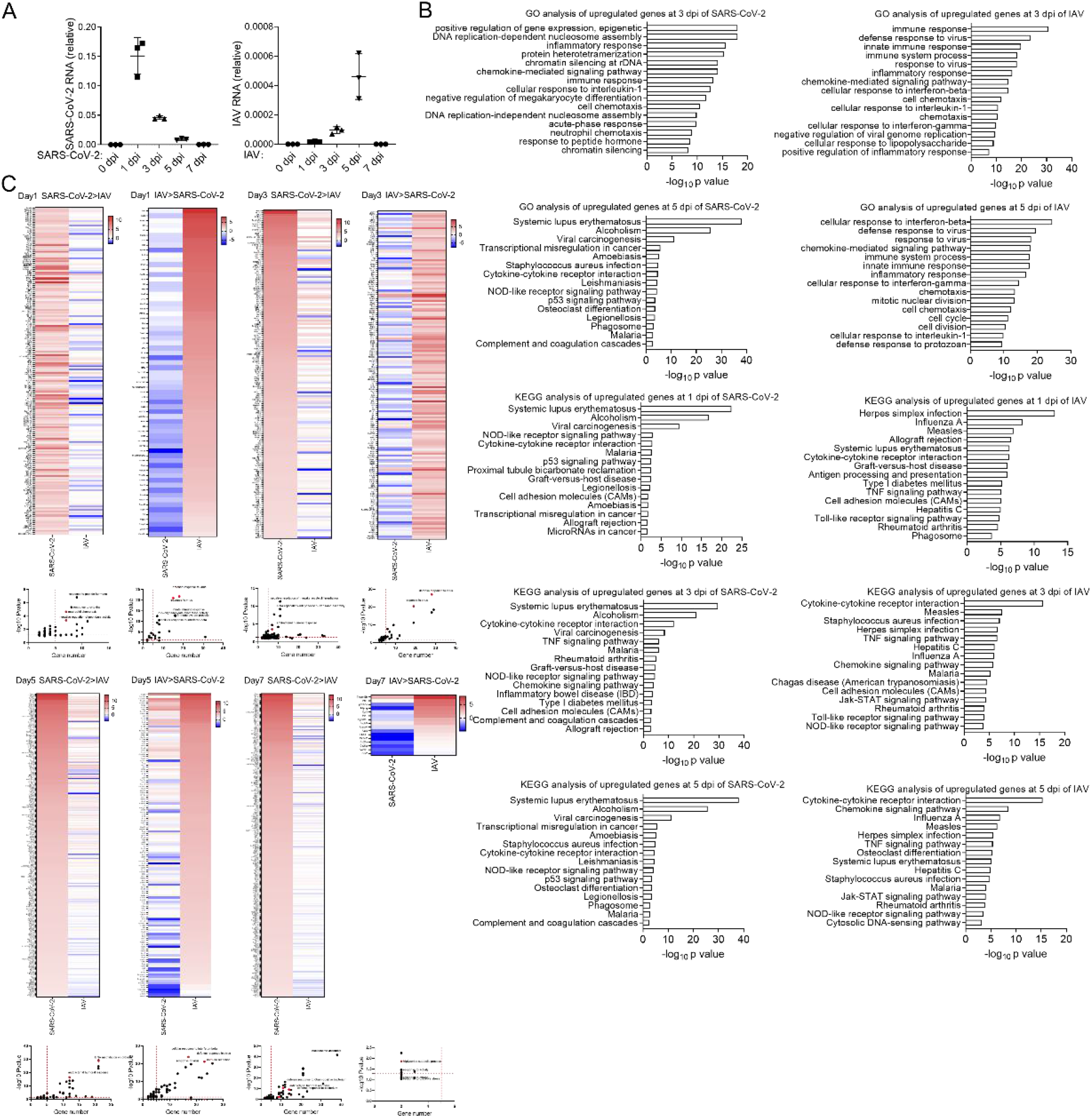
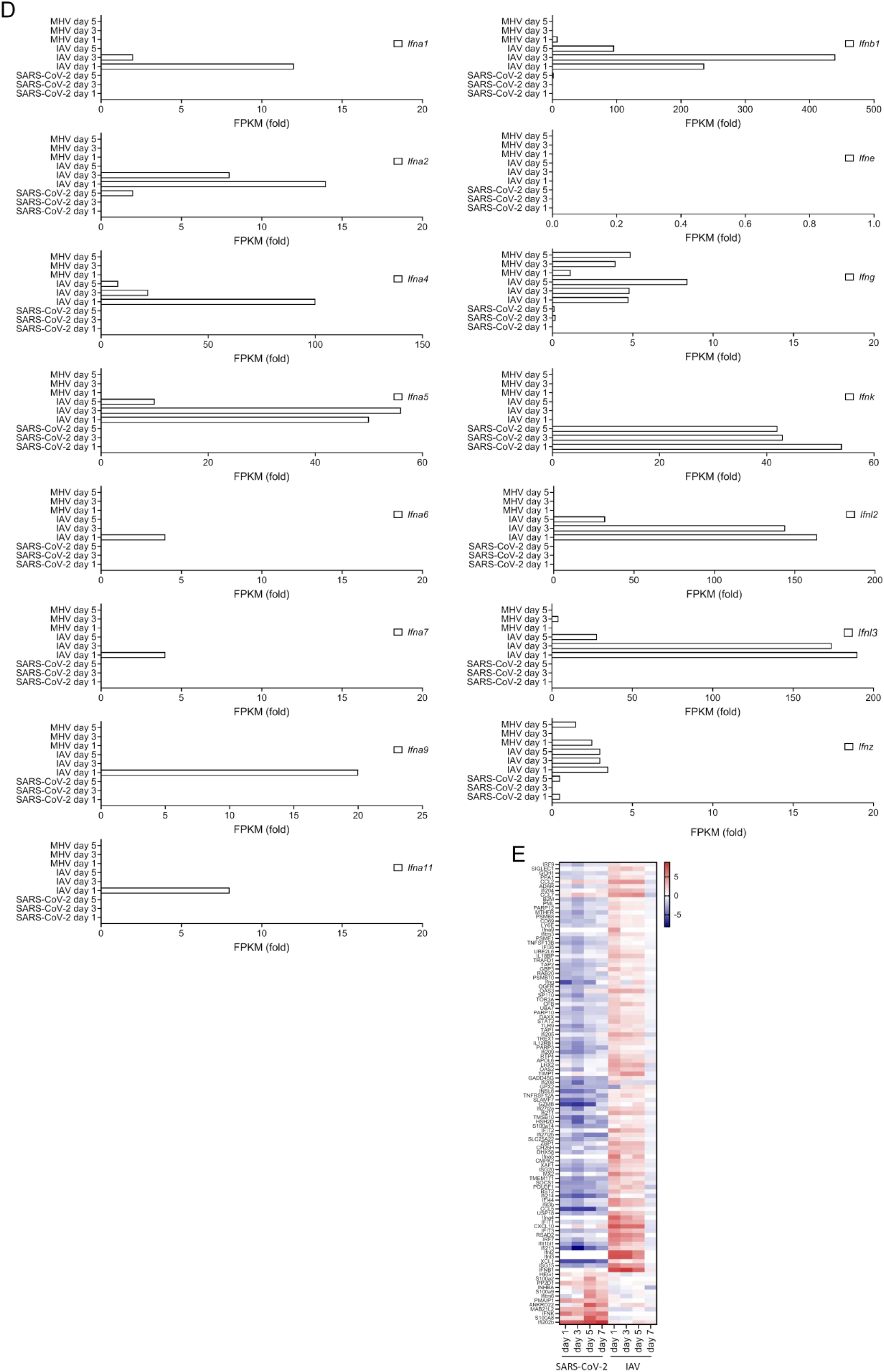
**(A)** qRT-PCR analysis for viral load in the lung of hACE2 mice infected intranasally with SARS-CoV-2 and C57BL/6 mice infected intranasally with IAV at 0 dpi, 1 dpi, 3 dpi, 5 dpi, and 7 dpi. *n = 3*. **(B)-(E)** RNA-seq analysis of RNA from lung of C57BL/6 mice infected with IAV and hACE2 mice infected with SARS-CoV-2 at 1dpi, 3 dpi, 5 dpi, and 7 dpi. Go and KEGG analysis was performed with the differentially expressed genes compared with Mock (B). Go and KEGG analysis was performed with the differentially expressed genes between IAV induced genes and SARS-CoV-2 induced genes (C) (*FC > 4 or < 0*.*25, P value < 0*.*05*). The expression of *IFNs* (D) and *ISGs* was analyzed (E).

**Figure S3.**
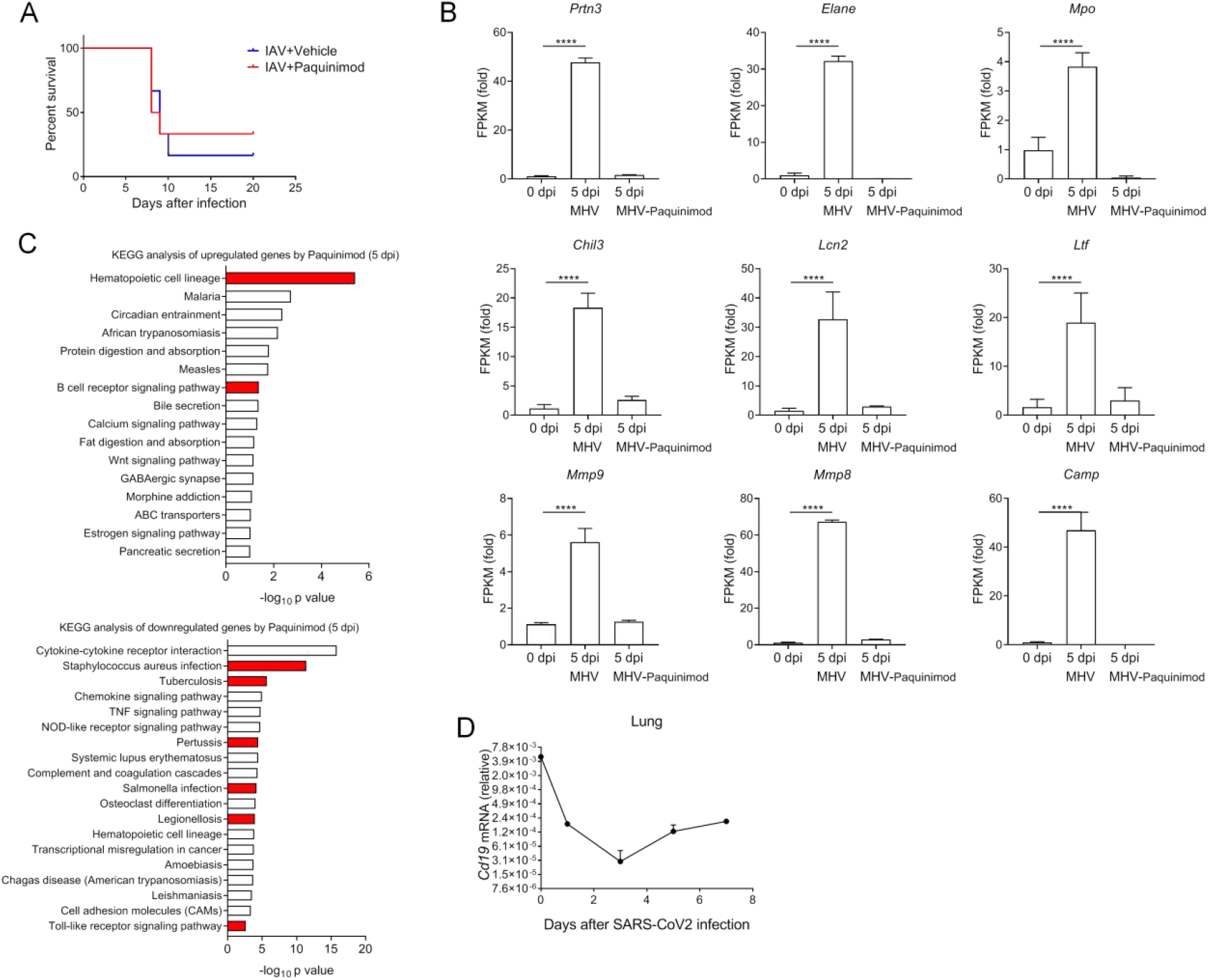
**(A)** Post-infection survival curves of C57BL/6 mice infected intranasally with IAV between Vehicle treated group and Paquinimod treated group. *n=6/group*. **(B)&(C)** RNA-seq analysis of RNA from lung of *Ifnar*^-/-^ mice infected intranasally with MHV between Vehicle treated group and Paquinimod treated group at 5 dpi. Analysis of the expression level of indicted genes (B). KEGG analysis was performed with the differentially expressed genes (C) (*FC > 4 or < 0*.*25, P value < 0*.*05*). **(D)** qRT-PCR analysis for the expression level of *Cd19* in the lungs of hACE2 mice infected intranasally with SARS-CoV-2 at 0 dpi, 1 dpi, 3 dpi, 5 dpi, and 7 dpi. *n = 3*. (*\*\*\*\*P < 0*.*0001*).

**Figure S4.**
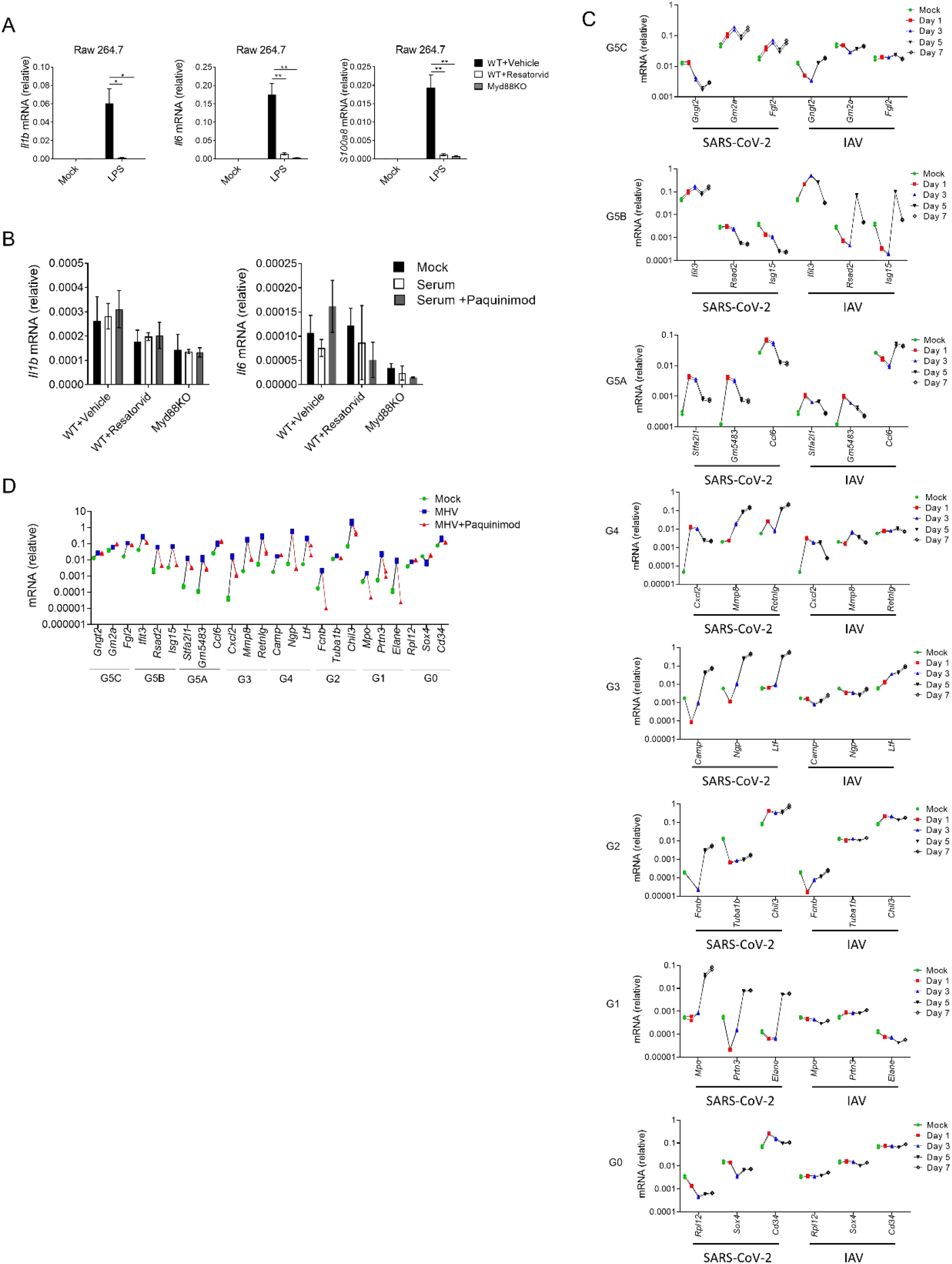
**(A)** qRT-PCR analysis for the expression level of *Il1b, Il6*, and *S100a8* in the Raw 264.7 cells stimulated with LPS among WT control group, WT Resatorvid treated group, and Myd88-/- group. *n = 3*. **(B)** qRT-PCR analysis for the expression level of Il1b and Il6 in the Raw 264.7 cells treated with indicated serum among WT control group, WT Resatorvid treated group and Myd88-/- group. *n = 3*. **(C)** RNA-seq analysis of the change of several related genes of neutrophil groups from the lungs of C57BL/6 mice infected with IAV and hACE2 mice infected with SARS-CoV-2 at 0 dpi, 1 dpi, 3 dpi, 5 dpi, and 7 dpi. **(D)** RNA-seq analysis of the change of several related genes of neutrophil groups from the lungs of Ifnar-/- mice infected intranasally with MHV between Vehicle treated group and Paquinimod treated group at 5 dpi. (*\*P < 0*.*05; **P < 0*.*01*).

**Table S1.**
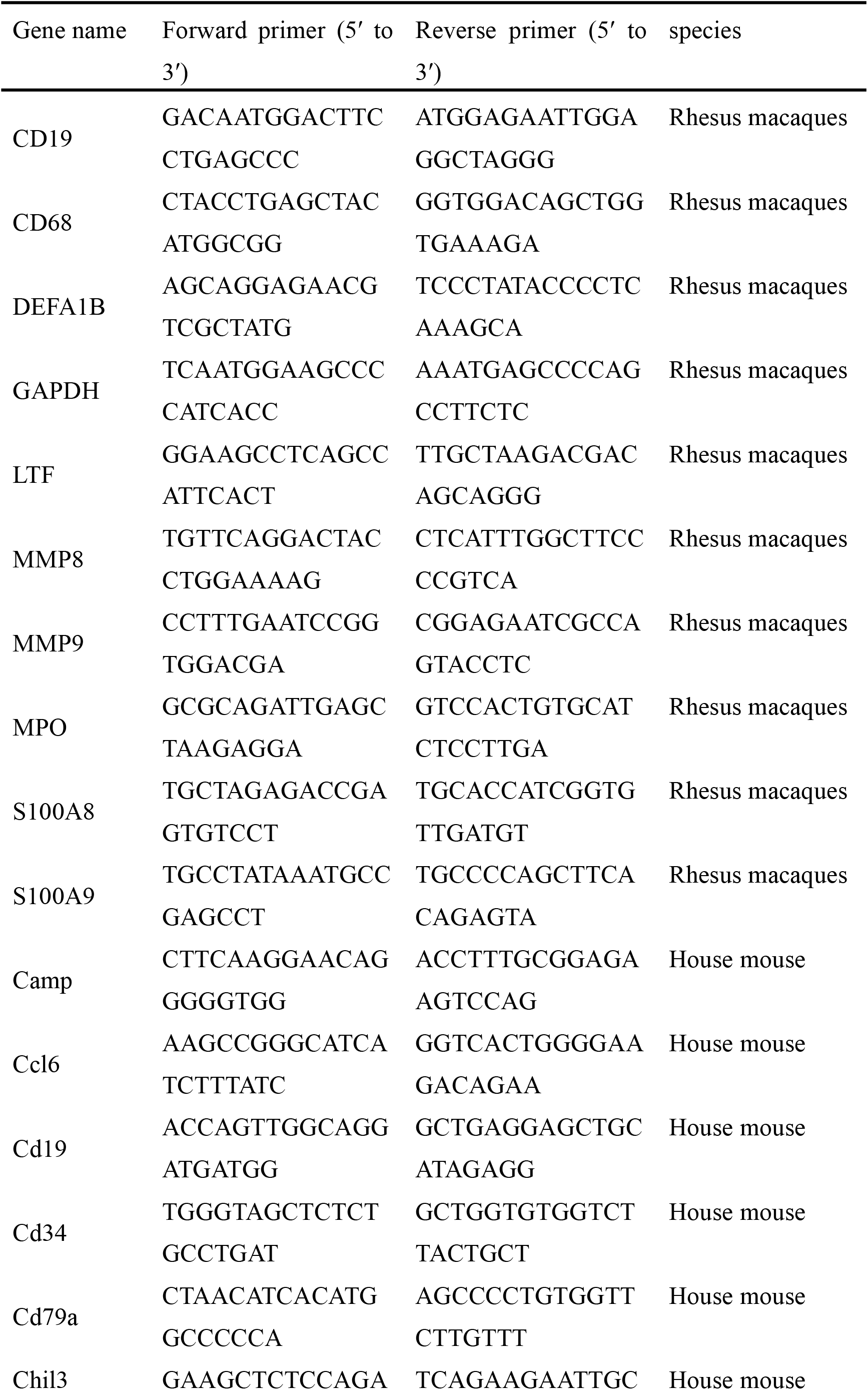
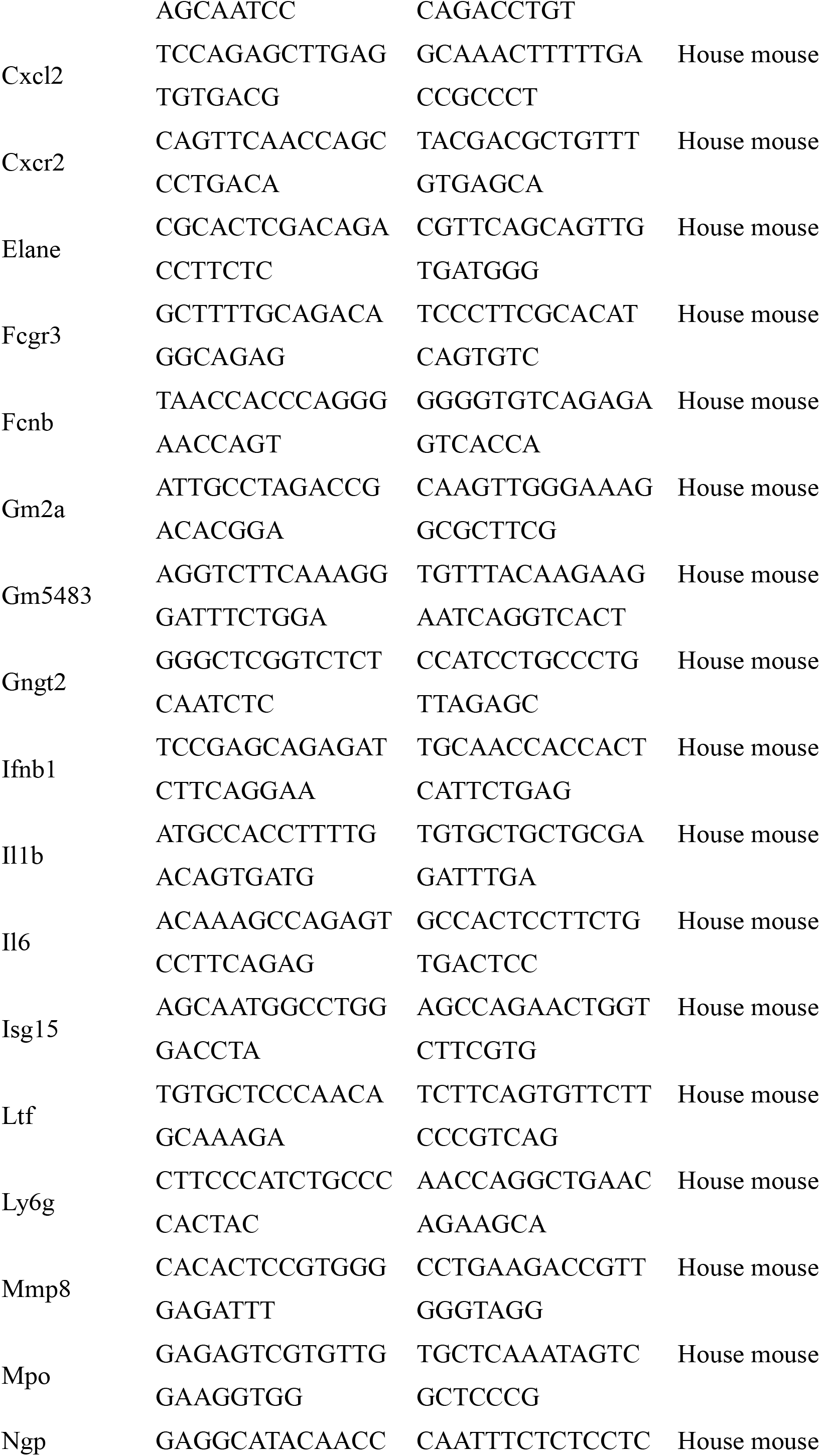
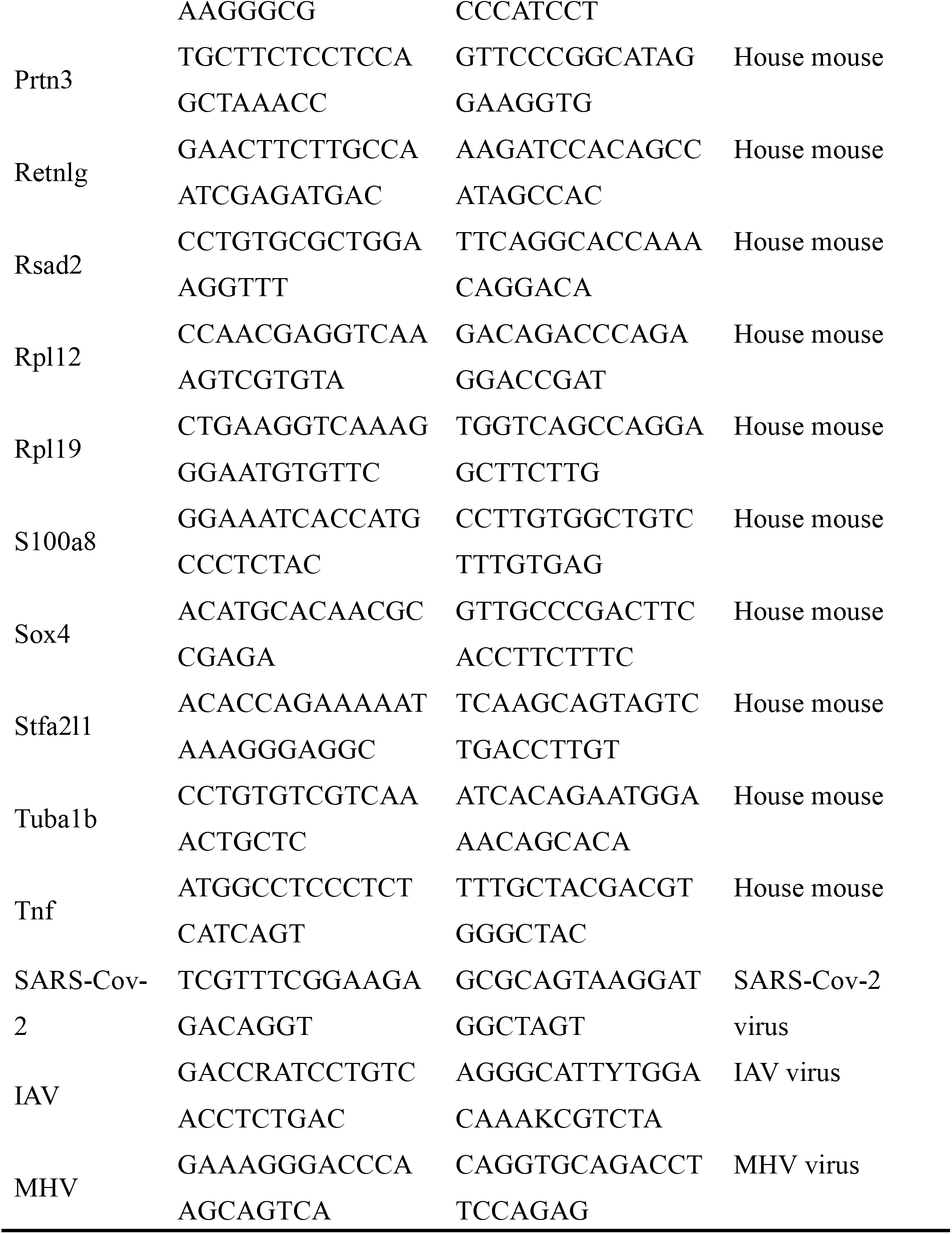
Sequences of qRT-PCR primers.

## References

Akira, S., Uematsu, S., and Takeuchi, O. (2006). Pathogen recognition and innate immunity. Cell 124, 783–801. doi: 10.1016/j.cell.2006.02.015

Bianchi, M.E. (2007). DAMPs, PAMPs and alarmins: all we need to know about danger. J Leukoc Biol 81, 1–5. doi: 10.1189/jlb.0306164

Bjork, P., Bjork, A., Vogl, T., Stenstrom, M., Liberg, D., Olsson, A., Roth, J., Ivars, F., and Leanderson, T. (2009). Identification of human S100A9 as a novel target for treatment of autoimmune disease via binding to quinoline-3-carboxamides. PLoS Biol 7, e97. doi: 10.1371/journal.pbio.1000097

Blanco-Melo, D., Nilsson-Payant, B.E., Liu, W.C., Uhl, S., Hoagland, D., Moller, R., Jordan, T.X., Oishi, K., Panis, M., Sachs, D., et al. (2020). Imbalanced Host Response to SARS-CoV-2 Drives Development of COVID-19. Cell 181, 1036–1045 e1039. doi: 10.1016/j.cell.2020.04.026

Bost, P., Giladi, A., Liu, Y., Bendjelal, Y., Xu, G., David, E., Blecher-Gonen, R., Cohen, M., Medaglia, C., Li, H., et al. (2020). Host-Viral Infection Maps Reveal Signatures of Severe COVID-19 Patients. Cell 181, 1475–1488 e1412. doi: 10.1016/j.cell.2020.05.006

Chan, J.K., Roth, J., Oppenheim, J.J., Tracey, K.J., Vogl, T., Feldmann, M., Horwood, N., and Nanchahal, J. (2012). Alarmins: awaiting a clinical response. J Clin Invest 122, 2711–2719. doi: 10.1172/JCI62423

Chen, G.Y., and Nunez, G. (2010). Sterile inflammation: sensing and reacting to damage. Nat Rev Immunol 10, 826–837. doi: 10.1038/nri2873

Cher, J.Z.B., Akbar, M., Kitson, S., Crowe, L.A.N., Garcia-Melchor, E., Hannah, S.C., McLean, M., Fazzi, U.G., Kerr, S.C., Murrell, G.A.C., et al. (2018). Alarmins in Frozen Shoulder: A Molecular Association Between Inflammation and Pain. Am J Sports Med 46, 671–678. doi: 10.1177/0363546517741127

Deng, Q., Sarris, M., Bennin, D.A., Green, J.M., Herbomel, P., and Huttenlocher, A. (2013). Localized bacterial infection induces systemic activation of neutrophils through Cxcr2 signaling in zebrafish. J Leukoc Biol 93, 761–769. doi: 10.1189/jlb.1012534

Diao, B., Wang, C., Tan, Y., Chen, X., Liu, Y., Ning, L., Chen, L., Li, M., Liu, Y., Wang, G., et al. (2020). Reduction and Functional Exhaustion of T Cells in Patients With Coronavirus Disease 2019 (COVID-19). Front Immunol 11, 827. doi: 10.3389/fimmu.2020.00827

Foell, D., Frosch, M., Sorg, C., and Roth, J. (2004). Phagocyte-specific calcium-binding S100 proteins as clinical laboratory markers of inflammation. Clin Chim Acta 344, 37–51. doi: 10.1016/j.cccn.2004.02.023

Giri, K., Pabelick, C.M., Mukherjee, P., and Prakash, Y.S. (2016). Hepatoma derived growth factor (HDGF) dynamics in ovarian cancer cells. Apoptosis 21, 329–339. doi: 10.1007/s10495-015-1200-7

Hadjadj, J., Yatim, N., Barnabei, L., Corneau, A., Boussier, J., Smith, N., Pere, H., Charbit, B., Bondet, V., Chenevier-Gobeaux, C., et al. (2020). Impaired type I interferon activity and inflammatory responses in severe COVID-19 patients. Science 369, 718–724. doi: 10.1126/science.abc6027

Huang, C., Wang, Y., Li, X., Ren, L., Zhao, J., Hu, Y., Zhang, L., Fan, G., Xu, J., Gu, X., et al. (2020). Clinical features of patients infected with 2019 novel coronavirus in Wuhan, China. The Lancet 395, 497–506. doi: 10.1016/s0140-6736(20)30183-5

Kang, R., Chen, R., Zhang, Q., Hou, W., Wu, S., Cao, L., Huang, J., Yu, Y., Fan, X.G., Yan, Z., et al. (2014). HMGB1 in health and disease. Mol Aspects Med 40, 1–116. doi: 10.1016/j.mam.2014.05.001

Kuri-Cervantes, L., Pampena, M.B., Meng, W., Rosenfeld, A.M., Ittner, C.A.G., Weisman, A.R., Agyekum, R.S., Mathew, D., Baxter, A.E., Vella, L.A., et al. (2020). Comprehensive mapping of immune perturbations associated with severe COVID-19. Sci Immunol 5. doi: 10.1126/sciimmunol.abd7114

Li, Y.M., Karlin, A., Loike, J.D., and Silverstein, S.C. (2002). A critical concentration of neutrophils is required for effective bacterial killing in suspension. P Natl Acad Sci USA 99, 8289–8294. doi: 10.1073/pnas.122244799

Liao, M., Liu, Y., Yuan, J., Wen, Y., Xu, G., Zhao, J., Cheng, L., Li, J., Wang, X., Wang, F., et al. (2020). Single-cell landscape of bronchoalveolar immune cells in patients with COVID-19. Nat Med 26, 842–844. doi: 10.1038/s41591-020-0901-9

Nathan, C. (2002). Points of control in inflammation. Nature 420, 846–852. doi: 10.1038/nature01320

Nauseef, W.M., and Borregaard, N. (2014). Neutrophils at work. Nat Immunol 15, 602–611. doi: 10.1038/ni.2921

Nicolas-Avila, J.A., Adrover, J.M., and Hidalgo, A. (2017). Neutrophils in Homeostasis, Immunity, and Cancer. Immunity 46, 15–28. doi: 10.1016/j.immuni.2016.12.012

Ometto, F., Friso, L., Astorri, D., Botsios, C., Raffeiner, B., Punzi, L., and Doria, A. (2017). Calprotectin in rheumatic diseases. Exp Biol Med (Maywood) 242, 859–873. doi: 10.1177/1535370216681551

Oppenheim, J.J., and Yang, D. (2005). Alarmins: chemotactic activators of immune responses. Curr Opin Immunol 17, 359–365. doi: 10.1016/j.coi.2005.06.002

Patel, S. (2018). Danger-Associated Molecular Patterns (DAMPs): the Derivatives and Triggers of Inflammation. Curr Allergy Asthma Rep 18, 63. doi: 10.1007/s11882-018-0817-3

Schelbergen, R.F., Geven, E.J., van den Bosch, M.H., Eriksson, H., Leanderson, T., Vogl, T., Roth, J., van de Loo, F.A., Koenders, M.I., van der Kraan, P.M., et al. (2015). Prophylactic treatment with S100A9 inhibitor paquinimod reduces pathology in experimental collagenase-induced osteoarthritis. Ann Rheum Dis 74, 2254–2258. doi: 10.1136/annrheumdis-2014-206517

Tan, L., Wang, Q., Zhang, D., Ding, J., Huang, Q., Tang, Y.Q., Wang, Q., and Miao, H. (2020). Lymphopenia predicts disease severity of COVID-19: a descriptive and predictive study. Signal Transduct Target Ther 5, 33. doi: 10.1038/s41392-020-0148-4

Vogl, T., Tenbrock, K., Ludwig, S., Leukert, N., Ehrhardt, C., van Zoelen, M.A., Nacken, W., Foell, D., van der Poll, T., Sorg, C., et al. (2007). Mrp8 and Mrp14 are endogenous activators of Toll-like receptor 4, promoting lethal, endotoxin-induced shock. Nat Med 13, 1042–1049. doi: 10.1038/nm1638

Wang, H., Liao, H., Ochani, M., Justiniani, M., Lin, X., Yang, L., Al-Abed, Y., Wang, H., Metz, C., Miller, E.J., et al. (2004). Cholinergic agonists inhibit HMGB1 release and improve survival in experimental sepsis. Nat Med 10, 1216–1221. doi: 10.1038/nm1124

Wang, S., Song, R., Wang, Z., Jing, Z., Wang, S., and Ma, J. (2018). S100A8/A9 in Inflammation. Front Immunol 9, 1298. doi: 10.3389/fimmu.2018.01298

Wilk, A.J., Rustagi, A., Zhao, N.Q., Roque, J., Martinez-Colon, G.J., McKechnie, J.L., Ivison, G.T., Ranganath, T., Vergara, R., Hollis, T., et al. (2020). A single-cell atlas of the peripheral immune response in patients with severe COVID-19. Nat Med 26, 1070–1076. doi: 10.1038/s41591-020-0944-y

Wu, C., Chen, X., Cai, Y., Xia, J., Zhou, X., Xu, S., Huang, H., Zhang, L., Zhou, X., Du, C., et al. (2020a). Risk Factors Associated With Acute Respiratory Distress Syndrome and Death in Patients With Coronavirus Disease 2019 Pneumonia in Wuhan, China. JAMA Intern Med. doi: 10.1001/jamainternmed.2020.0994

Wu, F., Zhao, S., Yu, B., Chen, Y.M., Wang, W., Song, Z.G., Hu, Y., Tao, Z.W., Tian, J.H., Pei, Y.Y., et al. (2020b). A new coronavirus associated with human respiratory disease in China. Nature 579, 265–269. doi: 10.1038/s41586-020-2008-3

Xie, X., Shi, Q., Wu, P., Zhang, X., Kambara, H., Su, J., Yu, H., Park, S.Y., Guo, R., Ren, Q., et al. (2020). Single-cell transcriptome profiling reveals neutrophil heterogeneity in homeostasis and infection. Nat Immunol. doi: 10.1038/s41590-020-0736-z

Yang, Han Z., and Oppenheim, J.J. (2017). Alarmins and immunity. Immunol Rev 280, 41–56. doi: 10.1111/imr.12577

Yang, Z., Du, J., Chen, G., Zhao, J., Yang, X., Su, L., Cheng, G., and Tang, H. (2014). Coronavirus MHV-A59 infects the lung and causes severe pneumonia in C57BL/6 mice. Virologica Sinica 29, 393–402. doi: 10.1007/s12250-014-3530-y

Yu, P., Qi, F., Xu, Y., Li, F., Liu, P., Liu, J., Bao, L., Deng, W., Gao, H., Xiang, Z., et al. (2020). Age-related rhesus macaque models of COVID-19. Animal Models and Experimental Medicine 3, 93–97. doi: 10.1002/ame2.12108

Zhang, B., Zhou, X., Qiu, Y., Song, Y., Feng, F., Feng, J., Song, Q., Jia, Q., and Wang, J. (2020). Clinical characteristics of 82 cases of death from COVID-19. PLoS One 15, e0235458. doi: 10.1371/journal.pone.0235458

Zheng, H.Y., Zhang, M., Yang, C.X., Zhang, N., Wang, X.C., Yang, X.P., Dong, X.Q., and Zheng, Y.T. (2020a). Elevated exhaustion levels and reduced functional diversity of T cells in peripheral blood may predict severe progression in COVID-19 patients. Cell Mol Immunol 17, 541–543. doi: 10.1038/s41423-020-0401-3

Zheng, M., Gao, Y., Wang, G., Song, G., Liu, S., Sun, D., Xu, Y., and Tian, Z. (2020b). Functional exhaustion of antiviral lymphocytes in COVID-19 patients. Cell Mol Immunol 17, 533–535. doi: 10.1038/s41423-020-0402-2

Zhou, P., Liu, Z., Chen, Y., Xiao, Y., Huang, X., and Fan, X.G. (2020). Bacterial and fungal infections in COVID-19 patients: A matter of concern. Infect Control Hosp Epidemiol, 1–2. doi: 10.1017/ice.2020.156

Zhu, N., Zhang, D., Wang, W., Li, X., Yang, B., Song, J., Zhao, X., Huang, B., Shi, W., Lu, R., et al. (2020). A Novel Coronavirus from Patients with Pneumonia in China, 2019. N Engl J Med 382, 727–733. doi: 10.1056/NEJMoa2001017

